# Alveolar and airway progenitors converge on an epigenetically encoded profibrotic state

**DOI:** 10.64898/2026.11.08.765110

**Authors:** Jessica Liang, Xulun Zhang, Rachel Genova, Anuradha Basyal, Kendall Foster, Qinxue Shen, Christopher Ting, Haichun Pan, Robert P. Dickson, Nicole Falkowski, Joshua Franzen, Jason R. Spence, Tristan Frum, Cyrina Ostgaard, Cole Weber, Kristine Konopka, Rubin Tuder, Lars Knudsen, Lauren Ayers, Cheng-Yu Lee, Darrell N. Kotton, Rachel L. Zemans

## Abstract

During organ regeneration, epithelial transitional states arise from progenitors and instruct the mesenchyme to remodel the extracellular matrix. However, the identity of the progenitors, the mechanisms underlying this transition, and why these states persist in tissue fibrosis often remain elusive. Here, using lineage tracing in mice, we show that the profibrotic *KRT5−/KRT17+* state in pulmonary fibrosis arises convergently from alveolar type (AT) 2 and airway secretory cells. AT1 cells generate neither *KRT5−/KRT17+* nor AT2 cells. TGF-β/SMAD3 acts through KMT2A and p300 to epigenetically reprogram human AT2 cells into the *KRT5−/KRT17+* state. In human pulmonary fibrosis, *KRT5−/KRT17+* cells slough off, and transitional cells re-epithelialize the denuded basement membrane. Transcriptionally convergent states arise in injured kidney, skin, and intestine. This work reframes fibrogenesis as the accumulation of a conserved profibrotic transitional state via ongoing turnover, not persistence alone, and nominates the state transition and its epigenetic regulators as therapeutic targets.

## INTRODUCTION

During organ regeneration after injury, resident progenitors proliferate and differentiate to replace lost cells and restore normal tissue architecture(1). Severe or repetitive injury unleashes plasticity, with progenitors adopting states and fates not present during homeostasis. Moreover, aberrant regeneration can result in excess extracellular matrix (ECM) deposition and disruption of tissue architecture – that is, organ fibrosis(2).

The lung epithelium is maintained by regionally distinct progenitors: the airways by basal and secretory cells and the alveoli by alveolar type (AT) 2 cells, which proliferate and differentiate into AT1 cells(3–13). Human respiratory airway secretory cells (RASC)(14, 15) and murine bronchioalveolar stem cells (BASCs)(16) localize to the distal airway-alveolar junctions, co-express secretory and AT2 cell markers, and generate both airway and alveolar progeny. After severe injury, secretory and basal cells mobilize to regenerate the alveolus(8, 17–31). Whether AT1 cells, historically thought to be terminally differentiated, can function as progenitors remains controversial(32–46).

Idiopathic pulmonary fibrosis (IPF) is a progressive fibrotic lung disease that culminates in respiratory failure due to severe architectural distortion of the lung(47). Fibroblastic foci (FF) – dome-shaped aggregates of fibroblasts in nascent collagen overlain by an epithelium, the likely active site of fibrogenesis, honeycomb cysts (HC), and dense mature collagen replace functional alveoli(47–53). IPF arises from impaired epithelial regeneration following repetitive injury, which activates mesenchymal cells at the FF to deposit excessive ECM(52, 54–56). However, which epithelial progenitor(s) give rise to the FF and HC is unclear, underscoring a longstanding debate over the alveolar versus airway origins of IPF(23–25, 28, 52, 57–62). Moreover, the specific stage of regeneration at which these progenitor(s) fail, the molecular mechanisms underlying the regenerative defect, and the “repetitive” nature of the injury are poorly understood.

A seminal advance was the identification of the *KRT5−/KRT17+* or “aberrant basaloid” cell state in IPF(63–67). These cells activate the profibrotic processes implicated in IPF pathogenesis - ER stress, TGF-β activation, and senescence(58, 68, 69) - and express a subset of basal cell markers(64, 66). *KRT5−/KRT17+* cells colocalize with profibrotic fibroblasts at the FF(66, 70–72) and directly activate fibroblasts(73–76), and their abundance correlates with disease progression and survival(77–79). Moreover, a transcriptionally and functionally similar “transitional” state arises and causes fibrosis in mice (Table S1)(63, 65, 67, 75, 80–85). Taken together, these findings suggest that accumulation of *KRT5−/KRT17+* cells is a critical regenerative defect driving the pathogenesis of IPF.

However, the origin of *KRT5−/KRT17+* cells has not been examined *in vivo*, in large part because mice were assumed incapable of generating them(86). Murine transitional cells, which arise from AT2 and secretory cells(31, 63, 67, 80–82, 87), lack some *KRT5−/KRT17+* markers, particularly basal cell markers, and their precise relationship to human *KRT5−/KRT17+* cells has been uncertain. Similarly, the origin of the basal cells that presumably give rise to the pseudostratified HC(51, 53, 60), is unclear: lineage tracing in mice implicates preexisting basal and secretory cells(23–25, 27–29), whereas cultured human AT2 cells transdifferentiate into basal cells via the *KRT5−/KRT17+* state(86), raising the possibility that FF and HC represent a single pathogenic process originating from AT2 cells.

The mechanisms through which progenitors give rise to *KRT5−/KRT17+* cells and why these cells accumulate in IPF also remain poorly understood. TGF-β induces the transitional and *KRT5−/KRT17+* state(65, 73, 76), but the breadth of transcriptional reprogramming suggests epigenetic rewiring, which has not been defined. Because transitional cells differentiate into AT1 cells as fibrosis resolves in mice, the state has been regarded as an intermediate specific to AT2-to-AT1 cell differentiation. Accordingly, the prevailing explanation for the accumulation of *KRT5−/KRT17+* cells in IPF is impaired AT1 cell differentiation, leaving cells arrested or “stuck” in this profibrotic state(63, 65, 81, 88–91).

Here, using lineage tracing across multiple mouse models of lung injury, we showed that *KRT5−/KRT17+* cells arose convergently from AT2 and airway secretory cells, reconciling the controversy over the alveolar versus airway origins of IPF. AT1 cells generated neither *KRT5−/KRT17+* nor AT2 cells, and we did not observe AT2-derived basal cells. In human iPSC-derived (i)AT2 cells, TGF-β/SMAD3 induced KMT2A and p300 to deposit activating histone marks at *KRT5−/KRT17+* genes and reprogram AT2 cells into this state. In human IPF, *KRT5−/KRT17+* cells sloughed off into the airspaces, and denuded basement membranes were re-epithelialized by transitional cells, suggesting ongoing turnover rather than, or in addition to, arrest underlies *KRT5−/KRT17+* cell accumulation, thereby nominating the progenitor-to-*KRT5−/KRT17+* cell transition and its epigenetic regulators as therapeutic targets. Finally, transcriptionally and functionally convergent states arose from epithelial progenitors in the injured kidney, skin, and intestine(92–96). Together, these findings identify the transitional state not as an intermediate of a single lineage, pathogenic only when cells arrest in it, but as a conserved epithelial progenitor stress response.

## RESULTS

### Diverse epithelial cell types intermingle at the FF, HC, and thickened alveolar septa in IPF

In IPF, FF, HC, and dense collagen replace healthy alveoli (Figure 1A), leaving vast areas devoid of alveolar epithelial cells (AECs) (Figure S1A). Spatial proximity has been taken to imply lineage relationships between cells in the FF and HC(86, 97, 98). Although spatial proximity cannot definitively establish lineage, we asked whether proximity is even suggestive — whether the juxtaposition of different epithelial cell types within a lesion is consistent and specific enough to nominate a progenitor. To test this, we characterized the epithelial cell composition of FF and HC across extensive areas of IPF tissue, rather than selected fields. However, we observed diverse combinations of epithelial cell types in close proximity at the FF, HC, and thickened septa. FF were typically covered by a monolayer of KRT5-KRT17*+* proSPC-cells, which also highly expressed additional markers of the *KRT5−/KRT17+* state such as KRT8 and p16 (Figures 1B, 1C, and S1B-S1E), consistent with prior reports(65, 66, 70, 72). Typically adjacent to the *KRT5−/KRT17+* cells at the FF were proSPC+ and/or SCGB3A2+ cells, which occasionally also overlaid the FF themselves (Figures 1D, 1E, S1F, and S1G). However, the *KRT5−/KRT17+* cells overlying FF were occasionally continuous with pseudostratified airways (Figures 1F and S1H), and rarely, KRT5+ KRT17+ basal cells overlaid the FF (Figures 1G and S1I).

**Figure 1.**
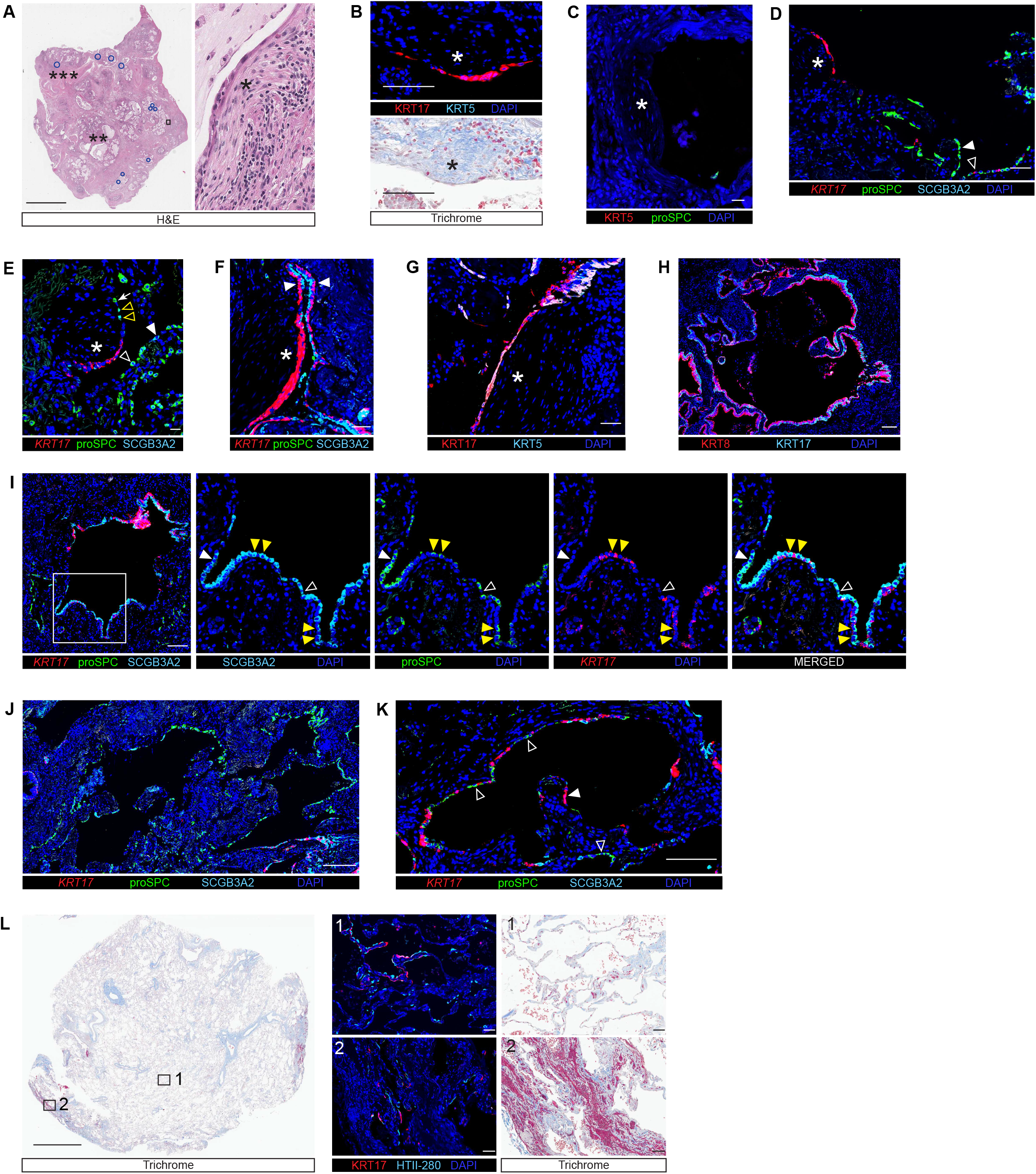
IPF lung is characterized by a microanatomically and molecularly haphazard epithelium in a severely remodeled environment. (A-K) Human IPF or (L) “Early UIP” lung stained with H&E, trichrome, or indicated antibodies and FISH probes. FF (*), HC (**), dense collagen (***). (A) All FF on section indicated by blue circles. (D) proSPC+ SCGB3A2-monolayer (closed arrowhead), *KRT17+* cells (open arrowhead). (E) proSPC+ SCGB3A2+ (open yellow arrowheads) and proSPC+ SCGB3A2-(white arrow) cells on FF. proSPC+ SCGB3A2-(open white arrowhead) and proSPC-SCGB3A2+ (closed white arrowhead) cells. (F) Pseudostratified airways (arrowheads). (I) Monolayer of proSPC-SCGB3A2+ cells (open arrowhead) and proSPC+ SCGB3A2+ cells (closed white arrowhead). proSPC+ SCGB3A2+ cells in pseudostratified epithelium (closed yellow arrowheads). (K) proSPC+ SCGB3A2-cells (open arrowheads), *KRT17+* cells (closed arrowhead). (L) Insets: Septa (1) and small peripheral regions of fibrosis (2). Scale Bars: 4 mm (A,L), 100 µm (B,H,I,K), 20 µm (C,E), 50 µm (D,F,G,L), and 200 µm (J). n≥3 patients. See also Figure S1.

The HC, irregularly shaped cysts 500-2000 µm in diameter, were typically lined by a pseudostratified epithelium with a KRT5+ KRT17+ basal layer (Figures 1H and S1J)(53). The structural similarity between HC and normal airways, in which basal cells give rise to luminal cells, has led to the widespread view that basal cells are the origin of the HC(24, 52, 53, 59). HC were occasionally populated by proSPC+ and/or SCGB3A2+ cells(53, 60) (Figures 1I-K and S1L) and, rarely, KRT5-KRT17+ monolayers were found in the HC (Figure S1K). Similar heterogeneity was evident in the tiny (100-500 µm) spherical cysts and on thickened septa. These are typically characterized by AT2 or secretory cell hyperplasia (Figure S1M, S1P, and S1Q), believed to be an early regenerative response to injury. However, they occasionally contained *KRT17+* cells or, rarely, were composed exclusively of KRT5-KRT17+ cells or a hyperplastic KRT5+ KRT17+ epithelium consistent with “peribronchiolar metaplasia” (Figures S1N, S1O, and S1R)(47).

Given the haphazard cellular organization of end-stage IPF, we sought regions in which alveolar architecture was relatively preserved and the earliest *KRT5−/KRT17+* cells might be observed as they emerge from their progenitors. On relatively normal septa without hyperplasia, we observed isolated *KRT17*+ proSPC+ and *KRT17*+ proSPC-cells (Figure S1S); these cells were KRT5-(Figure S1T). Moreover, we obtained tissue from IPF patients that was grossly normal (Figure 1L), which we termed “early usual interstitial pneumonia (UIP).” KRT17+ cells were identified adjacent to AT2 cells on normal and mildly thickened alveolar septa and in small peripheral regions of scarring (Figure 1L). These observations are consistent with emergence of the *KRT5−/KRT17+* state from AT2 or secretory cells before overt remodeling.

In summary, *KRT5−/KRT17+* cells were typically continuous with AT2 and secretory cells at the FF and emerged in the peripheral early UIP lung, and HC were typically lined by a pseudostratified epithelium. However, spatial proximity alone cannot establish lineage. Moreover, the striking variety of epithelial cell types juxtaposed on and around the FF and HC – revealing profound cellular and molecular disorganization – would render any such inference from spatial proximity uninformative. Computational trajectory analysis(64) is only predictive. To determine cellular origin, definitive *in vivo* lineage tracing is required.

### *KRT5−/KRT17+* Cells Arise from AT2 and Secretory Cells

*KRT5−/KRT17+ Cells Arise from AT2 Cells.* The assumption that human but not murine AT2 cells can generate *KRT5−/KRT17+* cells stems in part from the failure of murine AT2 cells to do so under specific 3D culture conditions(86). However, the “AT1-like” cells that emerge from murine AT2 cells in 2D culture(35) are predominantly transitional cells and upregulate *KRT5−/KRT17+* markers (Figure S2A)(88). We previously identified a murine analog of the *KRT5−/KRT17+* state by integrating three mouse models of lung regeneration: unbiased clustering revealed two clusters expressing transitional state markers, clusters 1 and 7 (Figure S2B)(88). Cells in cluster 7 expressed markers of the human IPF *KRT5−/KRT17+* state that most murine transitional cells do not. They expressed the specific basal cell markers that human *KRT5−/KRT17+* cells express such as *Krt17* (but not *Krt5*) and more highly expressed markers of senescence and TGF-β activation. Conversely, the top genes that distinguished cluster 7 from cluster 1 were highly differentially expressed in IPF *KRT5−/KRT17+* cells, suggesting that the human *KRT5−/KRT17+* program is what drives cluster 7 to cluster separately(88). Here, we found that at lower clustering resolution, clusters 1 and 7 merged into a single cluster, indicating they represent two subsets of murine transitional cells (Figure S2C). Thus, a subset of murine transitional cells closely recapitulates the *KRT5−/KRT17+* state.

To gain insight into whether the murine *KRT5−/KRT17+* state arises in additional injury models and derives from AT2 cells, we transferred cluster labels from this integrated reference dataset onto scRNA-seq datasets of AT2 lineage-labeled cells from three additional murine models: bleomycin, repetitive bleomycin, and pneumonectomy(44, 82, 99) (Table S2). In every model, the cells mapped to cluster 7 expressed higher levels of *KRT5−/KRT17+*, senescence, and TGF-β markers, as well as the basal cell markers expressed by human IPF *KRT5−/KRT17+* cells, but not *Krt5*. They also expressed genes comprising a discriminative signature of human *KRT5−/KRT17+* markers that murine transitional cells do not generally express (Figures S2D-S2F; Table S3-S4). Finally, scTOP, a computational algorithm that scores transcriptome-wide similarity between cell types(100), assigned human *KRT5−/KRT17+* cells the highest similarity score to the cluster 7 cells (Figures S2D-S2F). Together, these data suggest that a rare subset of murine transitional cells, arising from AT2 cells across many models of lung regeneration, closely recapitulates the human IPF *KRT5−/KRT17+* state. Of note, the murine and human *KRT5−/KRT17+* states are not identical: murine *KRT5−/KRT17+* cells are *Trp63*−, whereas 37% of human *KRT5−/KRT17+* cells express *TP63*(64). Only a minority of the murine cells expressed *Krt17* (Figures S2D-S2F). Because cluster assignment integrates thousands of genes and is robust to transcript dropout, and ∼15% of human *KRT5−/KRT17*+ cells also lack *KRT17* expression(64, 66), we speculate that the entire cluster is the murine counterpart of the human state. However, for the lineage-tracing experiments below, we adopted the more conservative definition of KRT17 protein expression.

To decisively determine whether murine AT2 cells generate *KRT5−/KRT17+* cells *in vivo*, we subjected SftpcCreERT2;mTmG mice to the single bleomycin, butylated hydroxytoluene (BHT), and repetitive bleomycin models of lung injury. In all models, cells expressing KRT17 but not KRT5 protein arose from both *Sftpc* lineage (lin)+ and *Sftpc* lin-cells (Figures 2A-2D). These cells expressed other markers of the *KRT5−/KRT17+* state, such as *Sprr1a* (Figure S2G). AT2 cells were likely progenitors of the *Sftpc* lin+ *KRT5−/KRT17+* cells, as BASCs were largely unlabeled with our tamoxifen regimen (Figure S2H) and *Sftpc* lin+ *KRT5−/KRT17+* cells emerged at the lung periphery (Figure S2I). Thus, in mice, *KRT5−/KRT17+* cells arise from AT2 cells and *Sftpc* lin-progenitors.

**Figure 2.**
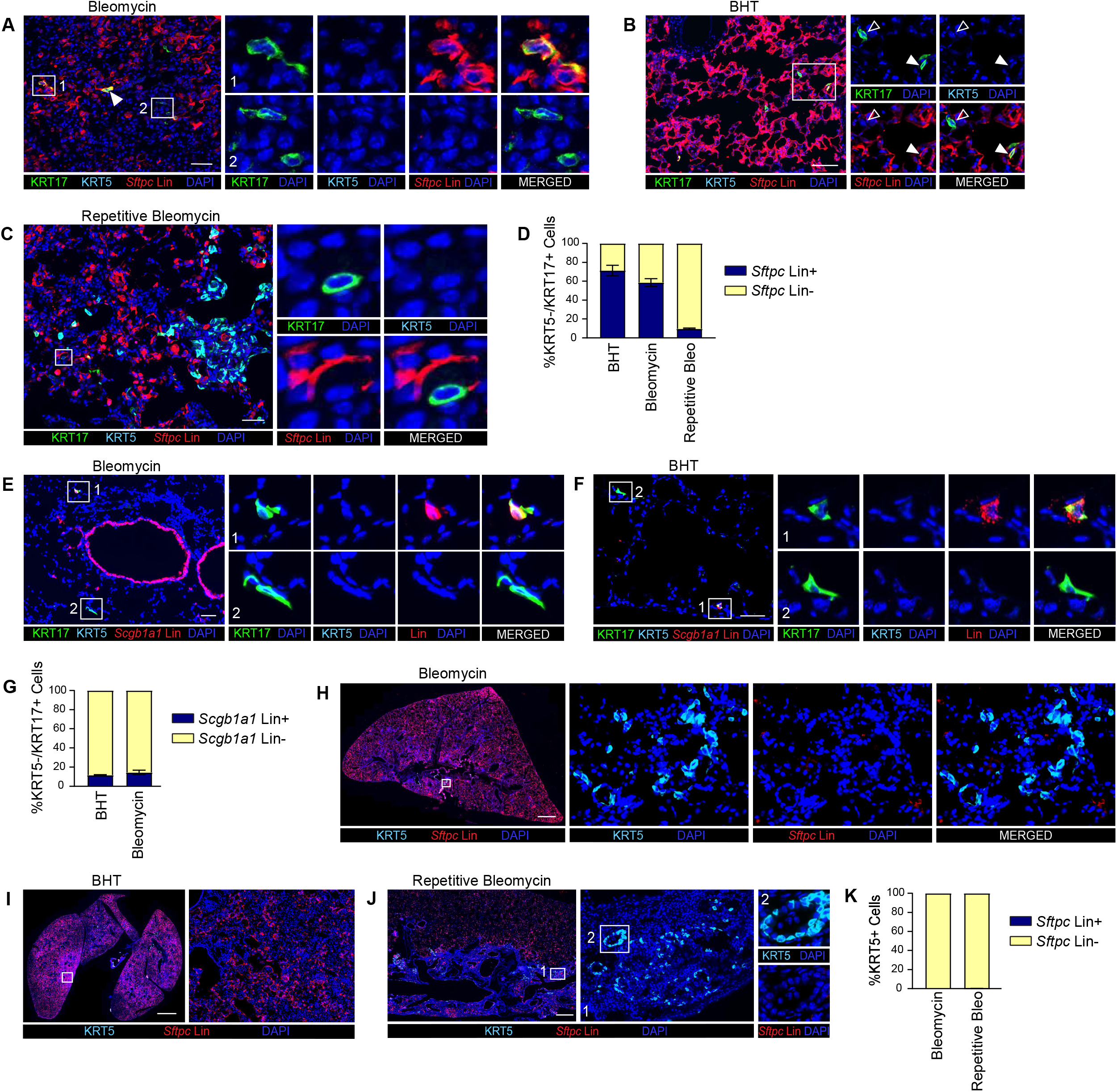
*KRT5−/KRT17+* cells arise from AT2 Cells. (A-D) *S*ftpcCreERT2;mTmG mice were treated with bleomycin (A), BHT (B), or repetitive doses of bleomycin (C). Lungs were immunostained for indicated antibodies. *Sftpc* lin+ cells (closed arrowheads). (D) Percent of KRT5-KRT17+ cells that were *Sftpc* lin+ or lin-. (E-G) *Scgb1a1CreERTM;tdTomato* mice were treated with bleomycin (E) or BHT (F). Lungs were stained with indicated antibodies. (G) Percent of KRT5-KRT17+ cells that were *Scgb1a1* lin+ or lin-. (H-K) *S*ftpcCreERT2;mTmG mice were treated with bleomycin (H), BHT (I), or repetitive bleomycin (J). Lungs were stained with indicated antibodies. (K) Percent of KRT5+ cells that were *Sftpc* lin+ or lin-. Scale Bars: 50 µm (A,E,F), 100 µm (B), 40 µm (C), 1 mm (H), 2 mm (I), and 500 µm (J). n=3-7 mice/group. Error bars represent Mean ± SEM. See also Figure S2.

*KRT5−/KRT17+ Cells Arise from Secretory Cells.* The *Sftpc* lin-progenitors of *KRT5−/KRT17+* cells may be secretory, basal, AT1, or (theoretically) other cells. Previous lineage tracing demonstrated that murine “transitional” cells arise from *Sox2*+/*Scgb1a1*+ progenitors(31, 63, 87), and computational analysis suggested that human *KRT5−/KRT17+* cells may arise from secretory cells in IPF(64). Therefore, we hypothesized that murine *KRT5−/KRT17+* cells arise from secretory cells *in vivo*. To test this hypothesis, we performed *Scgb1a1* lineage tracing in the single bleomycin and BHT models. *Scgb1a1* lin+ *KRT5−/KRT17+* cells were identified (Figures 2E-2G). Accordingly, *Sftpc* lin- and *Scgb1a1* lin+ *KRT5−/KRT17+* cells were peribronchiolar (Figures 2E and S2J). Together, these data suggest that the pathogenic *KRT5−/KRT17+* state arises convergently from AT2 and secretory progenitors.

### KRT5+ Basal Cells Do Not Arise from Sftpc Lin+ Progenitors

Although human AT2 cells transdifferentiate into basal cells *in vitro*(86), whether AT2 cells differentiate into basal cells *in vivo* is unclear. Basal cells arose in the distal lung, sometimes forming cystic structures, in the single(23, 28) and repetitive(101) bleomycin and (very rarely) BHT models; they were invariably *Sftpc* lin-(Figures 2H-2K and S2K). Interrogation of scRNA-seq datasets of *Sftpc* lineage tracing in multiple models confirmed that *KRT5*+ cells do not arise from AT2 cells (Figures S2D-S2F); this is consistent with prior reports showing that basal cells appearing in the distal lung after injury originate from airway basal and secretory cells(23–25, 28, 29). Assuming the luminal cells of the HC derive from basal cells, these data suggest that HC arise from airway, not AT2, cell progenitors.

### No evidence that AT1 Cells Differentiate into *KRT5−/KRT17+* or AT2 cells

Whether AT1 cells function as progenitors of AT2 cells during regeneration has been controversial(3, 32, 35, 37–39, 41, 43, 44, 46). AT1 lin+ cells acquire a transitional state gene signature during oncogenic KRAS-driven tumor progression(41, 42). However, whether AT1 cells give rise to the pathologic *KRT5−/KRT17+* cells in fibrosis has not been studied. To determine whether AT1 cells account for some of the *Sftpc* lin-progenitors of *KRT5−/KRT17+* cells and/or generate AT2 cells, we used two AT1 cell Cre drivers, *Rtkn2CreERT2* and *Igfbp2CreERT2,* which induce recombination in AT1, but not AT2, cells (Figures S3A and S3B). We observed no AT1 lin+ *KRT5−/KRT17+* cells in the bleomycin model (Figures 3A and 3B). In fact, we observed no AT1 lin+ AT2 cells (Figures 3C-3F) or any AT1 cell proliferation (Figures 3G-3J) in the hyperoxia, bleomycin, or repetitive bleomycin models. Repetitive bleomycin tested whether AT1 cells serve as a reserve progenitor pool recruited after resident AT2 progenitors are substantially depleted. Prior reports implicating AT1 cells as progenitors relied largely on *HopxCreER* mice(38, 39). *HopxCreER* labels some secretory cells (Figure S3C)(44), which are established AT2 cell progenitors(22), potentially reconciling our findings with the literature. Regardless, these data suggest that AT1 cells do not function as progenitors, consistent with prior reports(44, 46) but now established with independent Cre drivers, multiple injury models, and – for the first time to our knowledge – examining a *KRT5−/KRT17+* fate.

**Figure 3.**
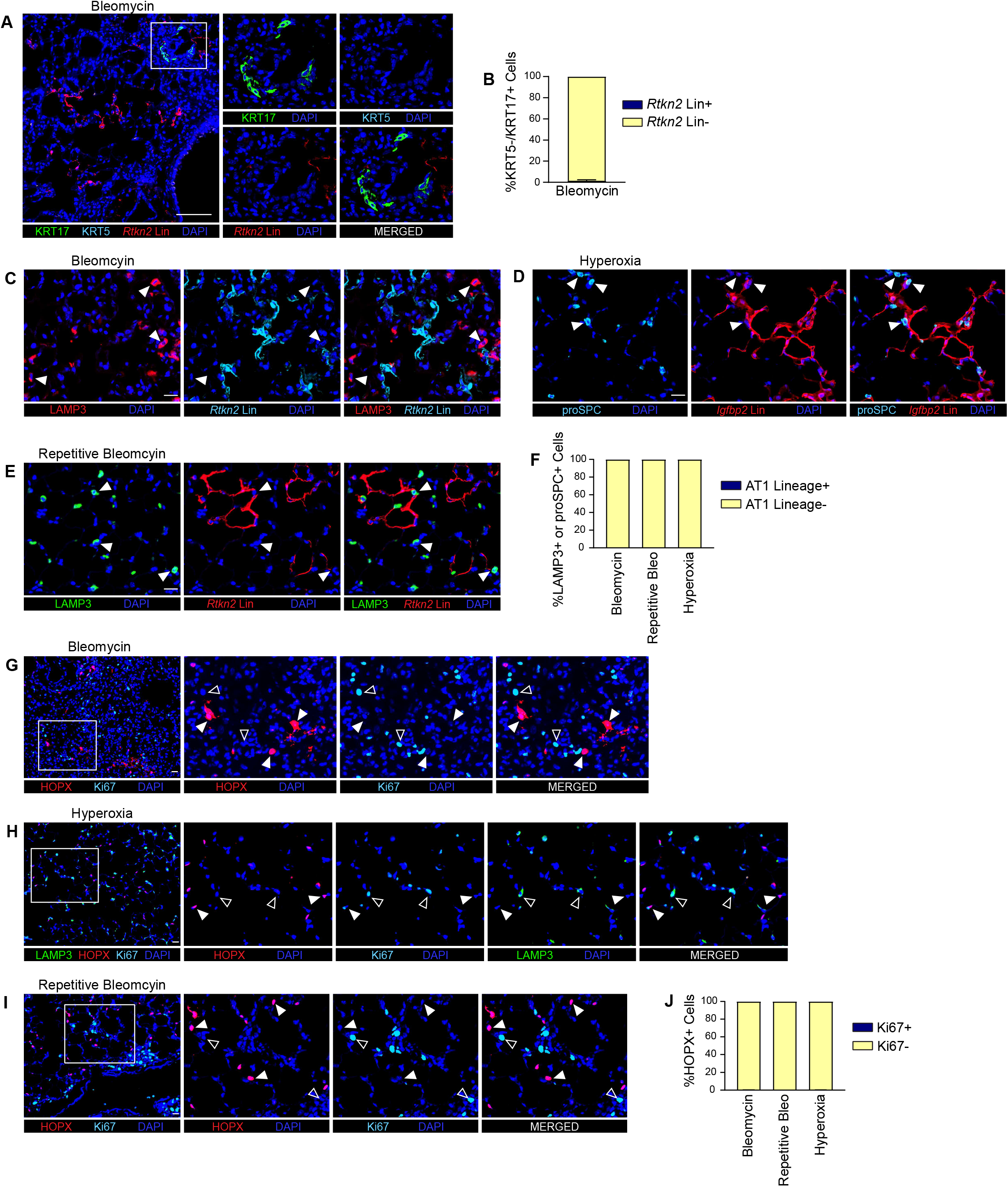
No evidence that AT1 Cells Differentiate into *KRT5−/KRT17+* or AT2 cells. *Rtkn2CreERT2;tdTomato* (A,C,E,G,I), *Igfbp2CreERT2;mTmG* (D), or *S*ftpcCreERT2;mTmG (H) mice were subjected to the bleomycin, hyperoxia, or repetitive bleomycin models of lung injury. Lungs were immunostained for indicated antibodies. (B) Quantitation of KRT5-KRT17+ cells that are *Rtkn2* lin+. (C-E) LAMP3+ or proSPC+ cells (arrowheads). (F) Quantitation of LAMP3+ or proSPC+ cells that are AT1 lin+. (G-I) HOPX+ cells (closed arrowheads) and Ki67+ cells (open arrowheads). (J) Quantitation of HOPX+ cells that are Ki67+. Scale Bars: 50 µm (A) and 20 µm (C-E,G-I). n=3-5 mice/group. Error bars represent Mean ± SEM. See also Figure S3.

### KMT2A and p300 are Necessary for TGF-β/SMAD3-Dependent AT2-to-*KRT5−/KRT17+* Cell Transition

*TGF-β/SMAD3 Induces the KRT5−/KRT17+ State.* Having established that AT2 cells are progenitors of *KRT5−/KRT17+* cells *in vivo*, we investigated the mechanisms regulating this transition. We previously demonstrated that TGF-β induces the transitional state in mouse and rat AT2 cells(65, 80). TGF-β also induced human iPSC-derived (i) AT2 cells to adopt the *KRT5−/KRT17+* state, as shown by upregulation of markers of transitional and basal cells and of the profibrotic processes active in this state (Figure S4A)(73, 74, 76, 86, 102). The effect was ALK5-dependent (Figure S4B;Table S5) and partially attenuated by factors that maintain AT2 identity (Figure S4C)(73). Basal cell markers remained lower than in mature basal cells (Figure S4D), indicating that TGF-β induces the *KRT5−/KRT17+*, but not basal cell, state. An unbiased survey of publicly available transcription-factor ChIP-seq datasets from lung epithelial cells identified SMAD3 as enriched at *KRT5−/KRT17+* marker genes (Figure S4E). TGF-β induced nuclear translocation of SMAD2/3 in iAT2 cells (Figure S4F), and SMAD3 inhibition attenuated the upregulation of *KRT5−/KRT17+* genes (Figure S4G;Table S5).

*KMT2A is Necessary for the AT2-to-KRT5−/KRT17+ State Transition.* Because the AT2 and *KRT5−/KRT17+* cell transcriptomes are markedly distinct and cell fate transitions are often epigenetically regulated, we hypothesized that this transition involves epigenetic reprogramming. The top 50 differentially expressed histone modifiers in *KRT5−/KRT17+* cells in IPF included *KMT2A* (Figure 4A)(64). We therefore stimulated AT2 cells with TGF-β in the presence of the KMT2A inhibitor MI-503. qPCR, western blotting, and RNA-seq confirmed that TGF-β upregulated *KRT5−/KRT17+* genes and KMT2A inhibition attenuated this response (Figures 4B-4E; Table S5). That KMT2A is necessary for the state transition was confirmed by CRISPR-mediated targeting of *KMT2A* (Figures 4F and S4H). Since KMT2A is an H3K4 trimethyltransferase, we performed H3K4me3 CUT&RUN on AT2 cells driven by TGF-β into the *KRT5−/KRT17+* state (Figure S4I and S4J). The 50 genes with the greatest H3K4me3 gain were highly upregulated in *KRT5−/KRT17+* cells in IPF (Figure 4G). Conversely, canonical *KRT5−/KRT17+* markers were significantly enriched for H3K4me3 following TGF-β (Figure 4H and S4K). Browser tracks illustrate H3K4me3 gain at *KRT5−/KRT17+* gene promoters, in intergenic regions, and diffusely across gene bodies (Figure 4I) — the latter a known feature of lineage-defining genes(103). Genes with greater H3K4me3 gain were more strongly inhibited by MI-503 (Figure 4J), suggesting KMT2A-dependent deposition drives *KRT5−/KRT17+* gene expression. Interestingly, some KMT2A-dependent genes showed no H3K4me3 gain (Figure S4L). This may reflect indirect regulation by KMT2A through intermediary factors; however, these genes tended to be enriched for H3K4me3 in AT2 cells (Figure S4M), suggesting KMT2A may be necessary to maintain this mark during the transition rather than establish it *de novo*. In fact, 87% of *KRT5−/KRT17+* genes were marked by H3K4me3 in AT2 cells (Figure S4K), suggesting that AT2 cells are epigenetically primed for this state transition. Together, these data suggest that KMT2A-mediated H3K4 trimethylation is essential for reprogramming AT2 cells into the *KRT5−/KRT17+* state.

**Figure 4.**
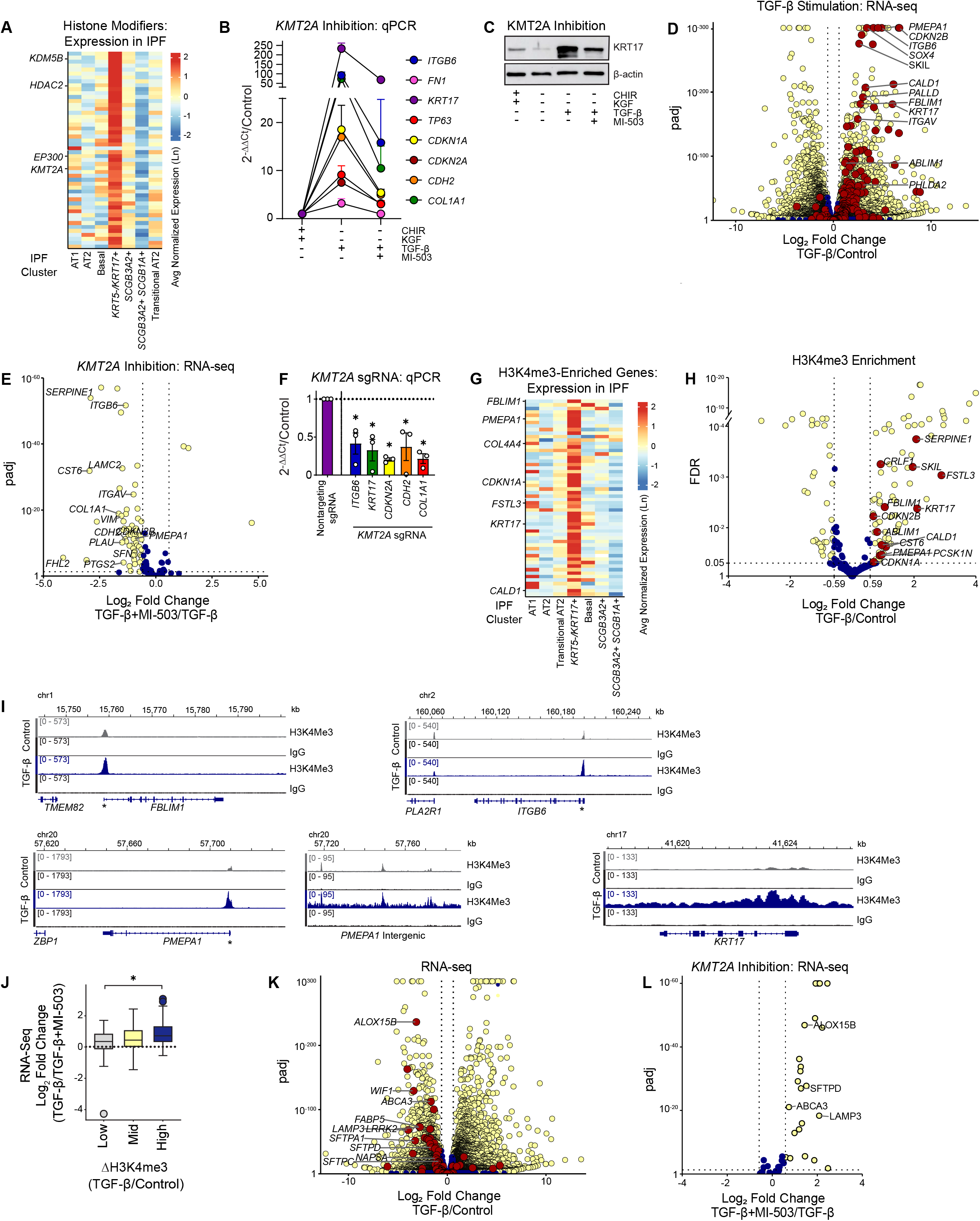
TGF-β/SMAD3 induces the *KRT5−/KRT17+* state via KMT2A-mediated H3K4 trimethylation. (A) Expression by the indicated cell types, as defined by human IPF scRNA-seq(64), of the top 50 differentially expressed (by FC) histone-modifying enzymes in *KRT5−/KRT17+* cells compared to all other epithelial cells. (B,C) Expression of indicated genes, as determined by qPCR (B) or western blot (C), in iAT2 cells treated with TGF-β ± MI-503. B) Statistics in Table S5. (D,E) RNA-seq of iAT2 cells treated with TGF-β ± MI-503. (D) All genes shown. (E) *KRT5−/KRT17+* marker genes statistically significantly upregulated in (D) shown. padj<0.05 by Wald test with Benjamini–Hochberg correction considered statistically significant. (F) qPCR for the indicated genes in non-targeting versus *KMT2A*-targeted iAT2 cells generated by lentiviral CRISPR-Cas9 and stimulated with TGF-β. *q<0.05 by one-sample t-test with Benjamini, Krieger and Yekutieli correction. (G) Expression by the indicated cell types in a human IPF scRNA-seq dataset(64) of the 50 genes with the greatest gain (by FC) of H3K4me3 in response to TGF-β, as determined by H3K4me3 CUT&RUN. (H) H3K4me3 marks in peaks annotated to protein coding genes in iAT2 cells treated with TGF-β compared to untreated cells, as determined by CUT&RUN. (I) Representative genome browser tracks showing mean H3K4me3 and IgG CUT&RUN signal at the indicated gene loci in untreated (Control) and TGF-β-treated iAT2 cells. * Transcription start site. (J) *KRT5−/KRT17+* marker genes upregulated by TGF-β (FC > 1.5, padj < .05) were ranked by magnitude of H3K4me3 gain and divided into tertiles. Box plots show the magnitude of MI-503–mediated inhibition, expressed as log₂FC (TGF-β/TGF-β+MI-503). Box, interquartile range; line, median; whiskers, 1.5 × IQR; circles, outliers. Kruskal-Wallis *H* = 7.13, *p* = .028; *p < .05 for low versus high tertile by Dunn test with Benjamini–Hochberg correction. (K) RNA-seq of iAT2 cells treated with or without TGF-β, all genes shown. Data also shown in D). (L) RNA-seq of iAT2 cells treated with or without TGF-β ± MI-503. Only AT2 genes downregulated (FC < -1.5, padj<0.05) in response to TGF-β shown. (D,E,H,K,L) Yellow dots: FC>1.5 or <0.67, FDR/pdj<0.05. Blue dots: FC 0.67-1.5 or FDR/padj>0.05. Red dots: *KRT5−/KRT17+* (D,H) or AT2 (K) marker genes. Dotted lines at p=0.05 and FC=1.5, 0.67. (B,F) Error bars: mean ± SEM. (B-F,H-L) n = 3 biological replicates. See also Figure S4.

TGF-β not only induced *KRT5−/KRT17+* genes but also suppressed AT2 identity genes (Figures 4K, S4N, and S4O). Conversely, the most strongly demethylated genes were highly expressed in AT2 cells (Figure S4P). MI-503 restored AT2 gene expression (Figure 4L), suggesting KMT2A suppresses the AT2 program during the *KRT5−/KRT17+* transition. Since H3K4me3 is an activating mark, this suppression is likely indirect. *ETV5*, a transcription factor maintaining AT2 identity(104), was suppressed by TGF-β, restored by MI-503, and depleted of H3K4me3 in response to TGF-β (Figure S4Q). *KDM5B*, a major H3K4me3 demethylase, was markedly upregulated in IPF *KRT5−/KRT17+* cells (Figure 4A) and is KMT2A-dependent (Figure S4R). Thus, KMT2A modestly silences the AT2 program, with KDM5B-mediated H3K4me3 demethylation at AT2 genes and their transcriptional regulators a possible mechanism.

*p300 is Necessary for the AT2-to-KRT5−/KRT17+ State Transition. EP300* (p300) is highly upregulated in the *KRT5−/KRT17+* state in human IPF (Figure 4A). Two mechanistically distinct p300 inhibitors attenuated the upregulation of *KRT5−/KRT17+* genes in response to TGF-β, as shown by qPCR, RNA-seq, and/or western blotting (Figures 5A-5D; Table S5)(74). Because p300 acetylates H3K27, typically at enhancers, we performed H3K27ac CUT&RUN (Figures S5A and S5B). The genes most highly enriched for H3K27ac after TGF-β were highly upregulated in the *KRT5−/KRT17+* state in IPF (Figure 5E). Conversely, *KRT5−/KRT17+* markers showed a broad gain in H3K27ac signal (Figures 5F, 5G, and S5C). 83% of *KRT5−/KRT17+* genes were marked by H3K27ac in AT2 cells (Figure S5C), suggesting that AT2 cells are epigenetically primed for this transition.

**Figure 5.**
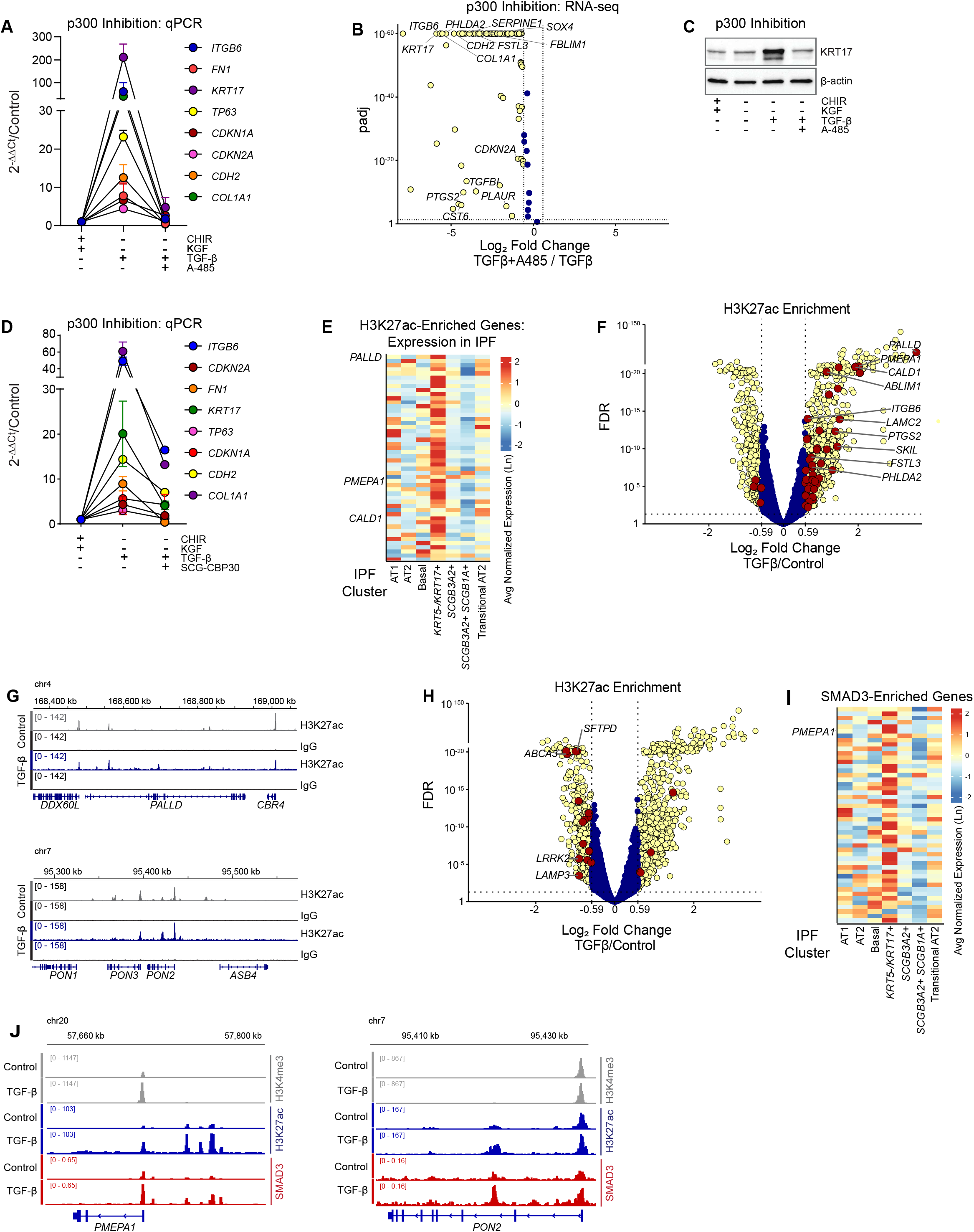
TGF-β/SMAD3 induces the *KRT5−/KRT17+* state via p300/CBP-mediated H3K27 acetylation. (A-D) Expression of indicated genes, as determined by qPCR (A,D), RNA-seq (B), or western blot (C), in iAT2 cells treated with TGF-β ± indicated inhibitors. (B) *KRT5−/KRT17+* genes shown. Yellow dots: |log₂FC| > 0.59, padj < 0.05. Blue dots: |log₂FC| < 0.59 or padj ≥ 0.05. Differential expression by DESeq2 Wald test with Benjamini–Hochberg correction. (C) Representative of 3 independent experiments. (A,D) Statistics in Table S5. (E-H) iAT2 cells were treated with TGF-β and subjected to H3K27ac CUT&RUN. (E) Expression by lung epithelial cell types in human IPF(64) of the 50 genes with the greatest gain of H3K27ac in response to TGF-β. (F,H) Differential H3K27ac signal in iAT2 cells treated with TGF-β compared to untreated (Control) cells. All peaks shown. Yellow dots: |log₂FC| > 0.59, FDR < 0.05, Red dots: *KRT5−/KRT17+* (F) or AT2 (H) cell marker genes |log₂FC| > 0.59, FDR < 0.05, Blue dots: |log₂FC| < 0.59 or FDR ≥ 0.05. DiffBind/DESeq2 with Benjamini–Hochberg correction. (G,J) Representative genome browser tracks showing mean H3K27ac (G) and SMAD3 (J) with IgG CUT&RUN signal at the indicated gene loci in untreated (Control) and TGF-β-treated cells. (B,F,H) Dotted lines at |log₂FC| = 0.59 and padj or FDR = 0.05. (I) Top 50 genes bound by SMAD3 in TGF-β-treated iAT2 cells, as determined by CUT&RUN. Expression in IPF(64) shown. (A-H) n=3 independent experiments. (I-J) n=2 independent experiments. See also Figure S5.

AT2 genes showed decreased H3K27ac (Figures 5H, S5D, and S5E), likely reflecting deacetylation, but p300 inhibition did not restore AT2 gene expression (Figure S5F), consistent with a prior report(74); further decreased expression may be due to limiting ongoing acetylation at AT2 genes amid a dynamic equilibrium. Notably, *KRT5−/KRT17+* marker genes were significantly enriched among genes downregulated by A-485 relative to all other tested protein-coding genes (79.4% vs. 21.7%; OR = 13.9, Fisher’s exact test, p = 5.4 × 10⁻^55^), and housekeeping genes were not substantially downregulated (Figure S5G), arguing against global transcriptional repression by p300 inhibition. Together, the functional requirement for p300 and the H3K27ac enrichment at *KRT5−/KRT17+* genes suggest that p300-mediated H3K27 acetylation is essential for *KRT5−/KRT17+* gene expression during the reprogramming of AT2 cells toward this state.

*KMT2A, p300, and SMAD3 Cooperate at KRT5−/KRT17+ Gene Loci via Distinct Mechanisms.* To gain insight into how KMT2A, p300, and SMAD3 interact at the chromatin, we performed SMAD3 CUT&RUN (Figures S5H and S5I). The genes most enriched for SMAD3 after TGF-β were highly upregulated in the *KRT5−/KRT17+* state in IPF (Figure 5I). Conversely, TGF-β increased SMAD3 occupancy at 70.5% of *KRT5−/KRT17+* genes, a significant enrichment compared with other genes (OR = 5.5, Fisher’s exact test, p = 3 × 10^⁻24^). At *KRT5−/KRT17+* loci, SMAD3 colocalized with H3K4me3 predominantly at promoters and with H3k27ac at both promoters and distal elements (Figure S5J). Representative tracks showing SMAD3, H3K4me3, and H3K27ac colocalization are shown in Fig. 5J. Together, these data suggest that SMAD3, KMT2A, and p300 act cooperatively to activate *KRT5−/KRT17+* genes.

### *KRT5−/KRT17+* Cells Undergo Turnover

Current dogma holds that the regenerative defect in IPF is epithelial cells “stuck” in the senescent *KRT5−/KRT17+* state due to impaired AT1 cell differentiation(63, 65, 81, 88–91). Though consistent, whether impaired differentiation is the cause of *KRT5−/KRT17+* cell accumulation and the paucity of AECs (Figure S1A) has not been proven. Like transitional cells(63, 88), *KRT5−/KRT17+* cells resolved in the bleomycin and BHT models of resolving fibrosis (though rare basal cells may remain) (Figures 6A and S6A). The density of transitional cells during fibrosis (Figure S6B), considered alongside the expansive morphology of AT1 cells, is difficult to reconcile with the universal differentiation of *KRT5−/KRT17+* cells into AT1 cells as fibrosis resolves. *KRT5−/KRT17+* cells express pro-apoptotic (Figure S6C) and anti-apoptotic(64) genes, the latter consistent with the apoptosis resistance of senescent cells. Finally, we previously observed *KRT5−/KRT17+* cytoplasmic proteins detectable in the circulation of IPF patients and KRT8^hi^ KRT17+ cells in mucus plugs in the airspace(78). Taken together, these observations raise the possibility that transitional and *KRT5−/KRT17+* cells resolve in part through cell death. We did not detect TUNEL+ *KRT5−/KRT17+* cells at FF (data not shown), perhaps reflecting the rapid clearance of apoptotic cells. However, we observed FF with *KRT5−/KRT17+* cells partially detached from the underlying basement membrane (Figure 6C), completely sloughed off (Figure 6D), and embedded in the airspace mucus plugs (Figures 6E and S6D). These observations are consistent with prior reports by us and others(70, 72, 78, 105, 106) and, since this tissue was immediately fixed after surgical biopsy, are unlikely to be artifact. Electron microscopy confirmed detachment of *KRT5−/KRT17+* cells, identified by the absence of lamellar bodies and prominent keratin filaments and desmosomes(97), from the underlying stroma (Figures 6F and 6G). Moreover, we observed septa with denuded basement membranes forming “clefts” invaginating into the remodeled stroma (Figure 6H), indicative of alveolar collapse, which is consistent with historical reports(107–110). That *KRT5−/KRT17+* cells accumulate despite cell death implies ongoing replenishment from progenitors. In fact, the progenitors, AT2 and secretory cells, tended to localize to the margins of the FF (Figures 1E and 6C). Moreover, we identified transitional epithelial cells, identified by the lack of lamellar bodies, that had reepithelialized the denuded clefts, appearing to seal the entrance to the former airspace (Figure 6I)(107). Our findings support a model of *KRT5−/KRT17+* cell emergence from progenitors, activation of the underlying mesenchyme to deposit excessive ECM, *KRT5−/KRT17+* cell death, alveolar collapse, and re-epithelialization by newly converted progenitors. Repeated cycles of this process would provide a mechanism for the prevailing but enigmatic model of IPF progression, in which individual FF mature into dense fibrosis devoid of epithelium, with new FF arising in adjacent, structurally normal alveoli, thereby propagating fibrosis across the lung; such cycles would also account for the elusive “repetitive” injury on which that model depends.

**Figure 6.**
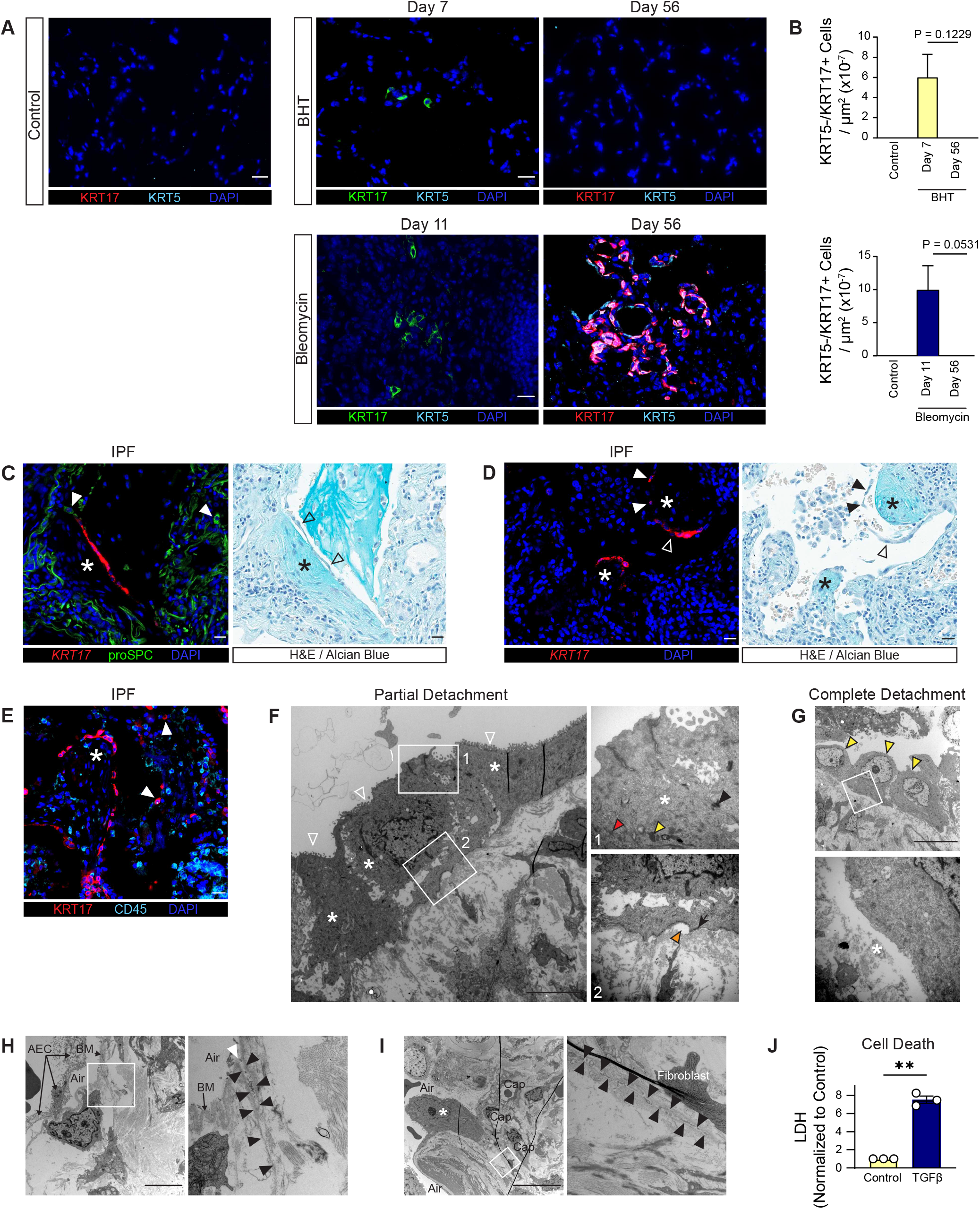
*KRT5−/KRT17+* Cells Undergo Turnover. (A,B) Mice were treated with BHT or bleomycin and lungs immunostained for indicated antibodies. (B) Quantitation of KRT5-KRT17+ cells at the indicated time points. Unpaired t-test with Welch’s correction. (C-E) IPF lungs were stained with H&E-Alcian Blue or indicated antibodies. FF (*). (C) Areas of cell detachment (open arrowheads) and proSPC+ cells on or near FF (closed arrowheads). (D) *KRT17+* cells detached from (open arrowheads) and sloughed off of (closed arrowheads) the FF. (E) KRT17+ CD45-cells in mucus plugs in airspaces (arrowheads). (F-I) IPF lungs were processed for transmission electron microscopy. (F) *KRT5−/KRT17+* cells (asterisks) with abundant thick bundles of intermediate filaments (red arrowhead), dense bodies (yellow arrowhead), desmosomes (black arrowhead), and hemidesmosomes (arrow), are partly detached (orange arrowhead) from the basement membrane. (G) *KRT5−/KRT17+* cells (yellow arrowheads) detached from the underlying stroma (inset). (H) Opening to slit-like cleft formed by apposed denuded basement membrane (white arrowhead) and walls of cleft (black arrowheads). Swollen alveolar epithelial cells detached from the basement membrane (AEC). BM: Basement membrane. (I) Cleft formed by apposed denuded basement membranes, invaginating deeply into the connective tissue (black arrowheads). A cuboidal epithelial cell (*) without lamellar bodies and with a prominent, spherical nucleolus, likely in transition from the AT2 to *KRT5-KRT17+* state, covering the opening to the collapsed cleft. Cap: Capillary lumen. (J) LDH measured in media of cultured *KRT5-/KRT17+* cells treated with TGF-β for 24 h. One sample t-test. **p<0.01. Scale Bars: 20 µm (A, C-E), 5 µm (F,H), 10 µm (G,I). n=3-6 mice/group (A,B), n≥3 patients (C-E), and n= 3 independent experiments (J). Error bars represent Mean ± SEM. See also Figure S6.

The mechanism of *KRT5−/KRT17+* cell death is unknown. Since we occasionally observed detached cells without morphologic features of apoptosis or necrosis (Figure 6F), suggesting that detachment may precede death, we wondered whether downregulation of cell-cell and cell-matrix adhesions, as has been reported in other epithelial cell types(111), may lead to anoikis. However, *KRT5−/KRT17+* cells upregulated cell-cell adhesion genes (Figure S6E); one might speculate that strong cell-cell adhesions contribute to the observed sloughing off of *KRT5−/KRT17+* cells in sheets (Figures 6C and 6D). Rather than loss of cell-matrix adhesions, there is a shift from the collagen IV-based alveolar basement membrane to a fibronectin/collagen I/III-rich provisional matrix and an associated upregulation of αvβ6/αvβ8 (Figure S6E)(48, 71). αvβ6 and αvβ8 activate latent TGF-β(69), resulting in high local concentrations of active TGF-β at the FF(112, 113). We found that TGF-β killed *KRT5−/KRT17+* cells in culture (Figure 6J). The mechanism leading to death and detachment of *KRT5−/KRT17+* cells at the FF, despite the resistance to apoptosis senescence confers, requires further study.

### The Transitional / KRT5−/KRT17+ State is a Stressed State Conserved Across Organs and Species

The transitional state was hypothesized to be a benign intermediate specific to AT2-to-AT1 cell differentiation(81). Subsequent work has shown that although transitional cells can differentiate into AT1 cells, they occupy a terminal position in trajectory analyses, resemble the pathogenic *KRT5−/KRT17+* state at the FF in IPF, and cause fibrosis in mice(75, 84, 85). They arise from multiple progenitors, in multiple species, in response to diverse stimuli and etiologies of lung injury/disease (Figures 1 and 2)(63, 67, 83, 114–117). This suggests that the state is not a benign intermediate specific to AT2-to-AT1 cell differentiation, but a conserved stress response of lung epithelial progenitors. We therefore asked whether a similar state arises from epithelial progenitors during injury and fibrosis in other organs. We interrogated scRNA-seq studies of murine and human skin, kidney, and intestinal epithelium, all of which have injury-emergent cell states(92–96).

In the murine kidney, a “damage-associated” proximal tubule (DA-PT) state emerges after injury (Figures 7A and S7A)(92). To determine whether it resembles the lung transitional and *KRT5−/KRT17+* states (hereafter, the “lung transitional states”), we generated gene modules representing the mouse and human lung transitional / *KRT5−/KRT17+* signatures; DA-PT cells scored highly for both (Figure 7B;Table S6), with upregulation of individual lung marker genes by the kidney DA-PT state shown in Figure S7B. Conversely, DA-PT marker genes were enriched in lung transitional / *KRT5−/KRT17+* cells in mouse models / human IPF (Figures 7C, S7C, and S7D;Table S6). By scTOP, DA-PT cells were most similar to the lung transitional state (Figure 7D), approaching the similarity scores observed between each lung state and either itself or the analogous state in other models and species (Figure S7E). Finally, the profibrotic pathways activated in the lung state - TGF-β, proteostasis, and senescence - were also activated in DA-PT cells (Figure 7E;Table S4). We applied these same analyses to the human kidney, mouse and human skin(95, 96), and mouse intestinal(94) epithelia (Figures 7F, 7K, 7T, S7F, S7I, and S7P). All behaved concordantly. The human adaptive proximal tubule (aPT) state (Figures 7G-J and S7G-S7H; Table S6)(93), the murine GA and human Bas-mig skin states (Figures 7L-7S and S7J-S7O;Table S6)(95, 96), and the murine intestinal SSC2 state (Figures 7U-7X and S7Q-S7S;Table S6)(94) each scored highly for the lung modules, showed reciprocal enrichment of their own markers in lung transitional / *KRT5−/KRT17+* cells, were most similar to the lung transitional / *KRT5−/KRT17+* states by scTOP, and activated the same profibrotic programs. The consistent emergence of a transcriptionally and functionally conserved state from diverse epithelial progenitors across organs, species, stimuli, and diseases identifies it as a fundamental, conserved epithelial progenitor stress response to injury.

**Figure 7.**
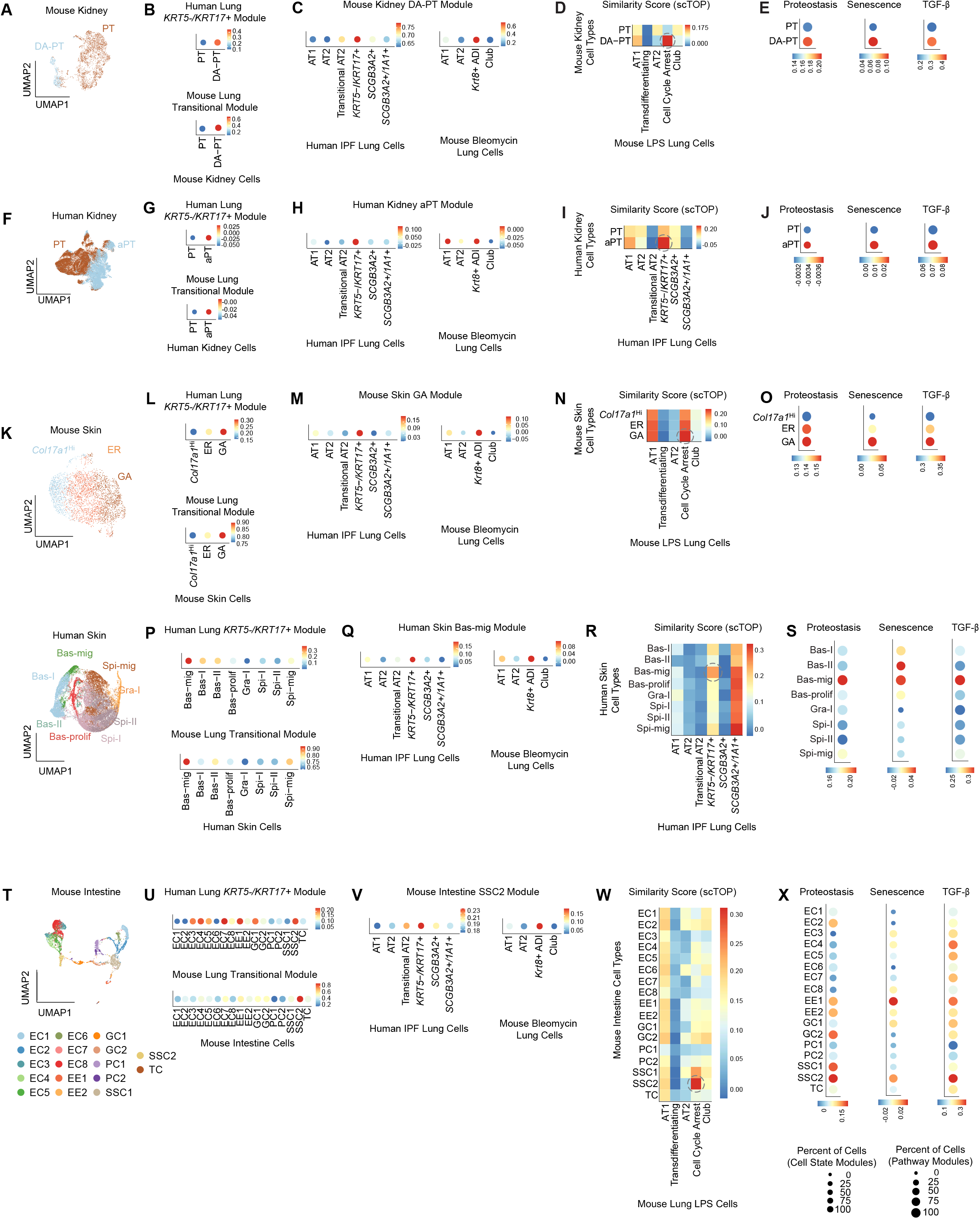
Conserved Cell States Emerge from Lung, Kidney, Skin, and Intestine. (A,F, K,T) UMAPs of injured (A) mouse(92) and (F) human(93) kidney proximal tubule (PT); (K) mouse(95) and human(96) skin; and (T) mouse small intestine(94). (B,G,L,P,U) Human lung *KRT5−/KRT17+* and mouse lung transitional state gene module activation by cells in (B) mouse and (G) human kidney PT cells, (L) mouse and (P) human skin basal cells, and (U) mouse intestinal epithelial cells. (C,H,M,Q,V) Gene module expression of the (C) mouse kidney DA-PT, (H) human kidney aPT, (M) mouse skin GA, (Q) human skin Bas-mig, and (V) mouse intestine SSC2 states by human and mouse lung epithelial cells(63, 64). (B,C,G,H,L,M,P,Q,U,V) Dot size represents percent of cells with a Cell State Module score > 0. (D,I,N,R,W) scTOP scores for (D) mouse and (I) human kidney PT, (N) mouse and (R) human skin, and (W) mouse intestine cell states compared with reference mouse(80) or human(64) lung cell types. Scale bars indicate the mean scTOP score. (E,J,O,S,X) Activation of gene modules of indicated pathways in the (E) mouse and (J) human PT, (O) mouse and (S) human skin, and (X) mouse intestine states. Dot size represents percent of cells with a Pathway Module score > 0. (B,C,E,G,H,J,L,M,O,P,Q,S,U,V,X) Scale bars indicate the Cell State or Pathway Module Score. See Supplemental Figure 7 for abbreviations. See also Figure S7.

## DISCUSSION

Impaired epithelial regeneration begets fibrosis in many organs, yet which progenitors are mobilized, the pathologic states they adopt, and the molecular programs that specify those fates remain critical unknowns. Here, we established the cellular origins of the profibrotic *KRT5−/KRT17+* cells in IPF *in vivo*. Lineage tracing demonstrated that *KRT5−/KRT17+* cells arise from AT2 and secretory cells, whereas AT2 cells did not generate basal cells. AT1 cells generated neither *KRT5−/KRT17+* nor AT2 cells. TGF-β directed SMAD3, KMT2A, and p300 to activate the *KRT5−/KRT17+* transcriptional program, establishing an epigenetic basis for this cell-fate decision. In human IPF, *KRT5−/KRT17+* cells detached and sloughed off into the airspaces, suggesting that their accumulation reflects dynamic turnover rather than only arrest. Together, these findings identify the progenitor-to-*KRT5−/KRT17+* cell transition, driven by epigenetic reprogramming, as the pivotal event in fibrogenesis (schematic).

### KRT5−/KRT17+ Cells Arise from AT2 and Secretory Cells

Although *KRT5−/KRT17+* cells are typically continuous with AT2 and secretory cells (Figure 1D) and the HC epithelium resembles and is continuous with pseudostratified airways(51, 53), the IPF epithelium is haphazard (Figures 1 and S1), and proximity cannot establish lineage. We therefore identified murine *KRT5−/KRT17+* cells and lineage traced their origin to AT2 and secretory progenitors (Figures 2 and S2)(88). These cells are rare in mice (Figure 2), as in human IPF (Figure 1A)(64, 66), but likely drive disease: *KRT5−/KRT17+* cells occupy the site of active fibrogenesis(66, 70, 71), activate TGF-β(69), directly activate fibroblasts(73, 74, 76, 118), cause fibrosis in mice(75, 84, 85), and predict disease progression in patients(77–79). That the same progenitors operate in human disease is supported by conserved progenitors across injury models, trajectory analyses in ferrets(117) and humans(64), activating histone marks at *KRT5−/KRT17+* loci in human AT2 cells (Figures S4K and S5C), the emergence of *KRT5−/KRT17+* cells on structurally normal septa in early disease (Figure 1L), and the *ex vivo* generation of *KRT5−/KRT17+* cells from human AT2 and secretory cells(73, 86, 102, 119). That murine AT2 cells generate *KRT5−/KRT17+* cells suggests mice and humans are more alike than assumed. Apparent species differences may reflect not intrinsic cell fate potential, but that our models – acute injury in young, isogenic, pathogen-free mice – do not unlock the plasticity observed in aged humans with genetic predispositions and environmental exposures. In other words, severity and chronicity unlock plasticity. Airway, not alveolar, progenitors generate basal cells in physiologic mouse models (Figures 2H-2K)(19, 23–25, 28, 59, 120), supporting an airway origin of HC. We cannot exclude an AT2 origin in IPF, however, since AT2 cells adopt a basal cell fate *in vitro*(86) and *in vivo* upon knockout of an epigenetic regulator required for AT2 identity(121), an extreme perturbation that may parallel the extreme disorganization of end-stage IPF; additionally, recent evidence suggests AT2 cells can give rise to secretory cells(122). Regardless, the dual alveolar and airway origins of *KRT5−/KRT17+* cells reconcile the longstanding debate over whether IPF is fundamentally an alveolar or a small-airways disease(61, 62).

### KRT5−/KRT17+ Cells Turn Over

The prevailing view holds that the pathologic *KRT5−/KRT17+* cells accumulate because they are arrested or “stuck” in this state due to impaired AT1 cell differentiation(63, 65, 81, 88–91); as a corollary, the therapeutic goal is to promote AT1 cell differentiation. We propose a paradigm shift: *KRT5−/KRT17+* cells are not simply stuck - they may differentiate into AT1 cells(74, 119) but they also die, as evidenced by the expression of cell death genes (Figure S6C), detachment from the underlying stroma (Figures 6C, 6F, and 6G), and sloughing into the airspaces (Figures 6D and 6E), leaving denuded alveolar septa (Figures 6C, 6D, 6H, and 6I). *KRT5−/KRT17+* cell death despite net accumulation implies continual re-emergence from progenitors, rather than only persistence of senescent cells that do not die or differentiate. Transitional cells reepithelialized the denuded septa (Figure 6I). Consistent with a pathogenic role for turnover, *KRT5−/KRT17+* cell proteins and transcripts in the blood and bronchioloalveolar lavage predict disease progression(78, 79). Thus, iterative cycles of *KRT5−/KRT17+* cell death and replenishment may drive fibrogenesis: progenitors at the margins of denuded septa (Figure 6C) assume the *KRT5−/KRT17+* state, activate the mesenchyme to deposit ECM at the FF, then die and are replenished(56).

How such cycles might be sustained remains to be determined. TGF-β/KMT2A/p300-dependent induction of αvβ6, which activates latent TGF-β, likely perpetuates this cycle by activating the ligand that recruits progenitors into the *KRT5−/KRT17+* state — and, to the extent that TGF-β also kills *KRT5−/KRT17+* cells (Figure 6J), couples recruitment to turnover. Regardless, this paradigm accords with the “spatial and temporal heterogeneity” of IPF(47, 50, 123) in which FF leave behind dense collagen devoid of epithelium and reappear on adjacent normal alveoli, advancing a “leading edge” of active fibrogenesis across the lung. It also reconciles how *KRT5−/KRT17+* cells, appearing rare in fixed tissue (Figure 1A), can drive widespread disease.

Epithelial cell death, alveolar collapse, and reepithelialization at the FF have been reported, but the dying cells were historically presumed to be resident AT2 cells(48, 50, 107, 109, 110, 124–132). This led to the dogma that AT2 cell death initiates IPF, and that preventing it is a therapeutic goal. We now recognize these cells as nascent repairing *KRT5−/KRT17+* cells. Therapy should therefore not necessarily aim to prevent AT2 cell death or promote the *KRT5−/KRT17+*-to-AT1 cell transition, which is futile once the septa are collapsed (Figures 6H and 6I), but at attenuating the progenitor-to-*KRT5−/KRT17+* cell transition.

### TGF-β/SMAD3 Induces KMT2A and p300 to Reprogram AT2 cells into the KRT5−/KRT17+ State

TGF-β activated KMT2A and p300 to coordinately deposit H3K4me3 and H3K27ac at many *KRT5−/KRT17+* genes (Figures 4, 5, S4, and S5). These loci are often marked with H3K4me3, H3K27ac, and SMAD3 in AT2 cells (Figures 4I, S4K, 5G, and S5C), suggesting the *KRT5−/KRT17+* program is epigenetically primed for rapid activation. This is consistent with evidence that the chromatin state of lung progenitors constraining their fates(133). KMT2A and p300 have been implicated in lineage specification across organs(134, 135), including p300 regulating *KRT5−/KRT17+* cell differentiation(74). Our findings also align with reports that loss of H3K27me3 drives the *KRT5−/KRT17+* transition(121, 136) – since H3K27 cannot carry both marks, demethylation frees the residue for p300-mediated acetylation. SMAD3 was required for the transition (Figure S4G) and colocalized with both marks (Figures 5J and S5J), consistent with the established cooperation of SMAD with KMT2A and p300(137). SMAD3 occupancy indicates that TGF-β activates *KRT5−/KRT17+* genes directly(138), beyond driving cell-cycle exit(65, 80). The transition requires repression of AT2 genes, which is also TGF-β-dependent (Figure 4K). AT2 genes lost H3K4me3 and H3K27ac (Figures S4N-S4P and S5D-S5F), and KMT2A inhibition partially restored their expression (Figure 4L). Because H3K4me3 is an activating mark, the silencing is likely indirect – plausibly eraser enzymes targeting regulators of AT2 identity (Figures 4A, S4Q, and S4R). Future work will elucidate the interactions between these and other(99, 139, 140) transcriptional regulators of this state transition. Interestingly, the mesenchymal marker expression and cell spreading previously interpreted as epithelial-mesenchymal transition (EMT)(22, 141, 142) likely reflect the AT2-to-*KRT5−/KRT17+* cell transition (Figure S4); hence, the mediators identified in the EMT literature are candidate regulators of *KRT5−/KRT17+* cells.

### No Evidence that AT1 Cells Function as Progenitors in Lung Regeneration

Whether AT1 cells are terminally differentiated remains controversial. An AT1 cell origin of *KRT5−/KRT17+* cells is plausible since AT1-derived cells express transitional state markers following oncogenic *Kras* expression(41, 42). We used two AT1-specific Cre drivers and/or injury models not previously applied to this question — including repetitive bleomycin, which extensively depletes AT2 cells, providing permissive conditions for AT1 progenitor function to emerge (Figure 3). AT1 cells did not generate *KRT5−/KRT17+* or AT2 cells - contrasting with some reports(38, 39, 43) but consistent with others(44–46). Earlier reports that “AT1-like” cells in 2D culture can differentiate into AT2 cells were interpreted as evidence of AT1 cell plasticity(34–37); we now know that these “AT1-like” cells are predominantly transitional cells(88), which can revert to AT2 cells(67, 74). AT1 cells may nonetheless act as progenitors in other contexts, such as ectopic expression of oncogenes or airway transcription factors(38, 39, 41–43, 143).

### The KRT5−/KRT17+ State is a Conserved Stressed State

The transitional/*KRT5−/KRT17+* state is a conserved response of diverse lung epithelial progenitors (Figure 2) to diverse stressors, across species, injury models and diseases, and lung development(41, 42, 63–67, 73, 80–83, 85, 114–117, 144–155). Even tracheal basal cells may adopt this state(91). That transcriptionally and functionally convergent states also emerge from epithelial progenitors in the kidney, skin, and intestine (Figure 7) establishes it as a conserved pan-epithelial progenitor stress response to injury. This contrasts with the view that the “transitional” state is a benign intermediate specific to AT2-to-AT1 cell differentiation, analogous to the graded intermediates of hematopoiesis and pathogenic only when cells arrest in it. Several observations argue otherwise: they express AT2 markers and *NKX2-1* at lower levels than AT1 cells(63, 64, 80); they occupy a terminal position on computational trajectories(63, 80, 88, 146, 156); they arise from secretory cells, they arise without AT1 cell death and therefore without a demand for AT1 cell regeneration; and they are profibrotic(75, 84, 85, 90). Moreover, a state defined by the AT2-to-AT1 cell trajectory cannot, by definition, arise in the kidney, skin, or intestine. Its identity is intrinsic - a stress program epithelial progenitors adopt upon injury, and the AT1 cell is one available fate, not its *raison d’être*. Genuine AT2-to-AT1 cell intermediates exist: the “transdifferentiating” or “AT2-AT1” cells identified in mice(80, 146, 155, 157) and the human “transitional” cells, which are transcriptionally more similar to AT1 than *KRT5−/KRT17+* cells (Figure S7E). *In vitro,* AT2 cells can differentiate into AT1 cells, bypassing the transitional state(155, 158); whether the transitional state is obligatory *in vivo* remains unknown.

The global transcriptional similarity and the engagement of the same fibrogenic programs — TGF-β, ER stress, senescence — suggest transitional cells may contribute to fibrogenesis in many organs. We speculate that this state evolved as an adaptive wound-healing mechanism(2, 127), transiently activating the mesenchyme to deposit matrix and restore barrier integrity during *acute* organ injury, but is hijacked into a pathologic self-amplifying loop in fibrosis. Whether the transcriptional and epigenetic regulators are conserved remains untested, but KMT2A and/or p300 inhibition attenuates fibrosis in the kidney, skin, and intestine(159–162). Targeting the transition of progenitor cells into this conserved state and its shared transcriptional and epigenetic regulators has the potential to treat impaired epithelial regeneration and fibrosis across organs.

## METHODS

### Mouse Models

Animal protocols were approved by the University of Michigan Institutional Animal Care Use Committee. SftpcCreERT2 mice were obtained from Dr. Harold Chapman (University of California San Francisco, San Francisco, CA)(163). *Rtkn2CreERT2* mice were obtained from Dr. Jichao Chen (University of Texas, Houston, TX)(164)*. Igfbp2CreERT2* mice were obtained from Dr. Nan Tang (National Institute of Biological Sciences, Beijing, China)(46) and were backcrossed to C57BL/6 mice for 10 generations. The following mice were purchased from The Jackson Laboratory: *Rosa26mT/mG* (#007576), *Rosa26tdTomato* (#007909), and *Scgb1a1CreERTM* (#016225). *S*ftpcCreERT2+/-;Rosa26mT/mG+/- (abbreviated *S*ftpcCreERT2;mTmG) and *Scgb1a1CreERTM^+/-^;Rosa26tdTomato^+/-^* (abbreviated *Scgb1a1CreERTM;tdTomato)* mice were fed tamoxifen chow (Inotiv TD.130859) for 2 w beginning at 6 w of age. *Rtkn2CreERT2^+/-^;Rosa26tdTomato^+/-^*(abbreviated *Rtkn2CreERT2;tdTomato)* mice were administered 4 mg tamoxifen (Sigma T5648) in corn oil i.p. every other day for 3 doses beginning at 4-6 w of age. *Igfbp2CreERT2^+/-^;Rosa26mT/mG^+/-^* (abbreviated *Igfbp2CreERT2;mTmG*) mice were administered 0.075 mg/g body weight tamoxifen i.p. every 24 h for a total of 5 doses beginning at 8 w of age. *HopxCreER^+/^;Rosa26mT/mG^+/-^*(abbreviated *HopxCreER;mTmG)* lung tissue was obtained from Drs. Tushar Desai and Nicholas Juul after administration of 2 doses of 4 mg tamoxifen i.p. at 6 months of age, with lungs harvested at 9 months of age. After a washout period of at least 3 weeks, mice were treated once with 0.75-1.8 U/kg bleomycin (Meitheal Pharmaceuticals NDC 71288-106-10, Fresenius Kabi C103610, or Hikma NDC-0143-9240-01) and euthanized 11, 14, 21, or 56 d post-injury, or treated with 1 U/kg bleomycin every other week for 3 doses and euthanized at 9 w after the final dose(101). Alternatively, mice were exposed to 95% oxygen for 3 d and euthanized after a 3-, 21- or 42-d recovery period in room air or were treated with 200-400 mg/kg BHT (Fisher Scientific S25212) and euthanized 5 or 7 d later. Lungs were fixed with 10% formalin (Epredia 5701) in 0.5% low melting point agarose (Promega V2111), processed, embedded in paraffin, and cut into 4 µM sections.

### Human IPF Tissue

The Department of Pathology at the University of Michigan database of formalin-fixed paraffin-embedded lung tissue was queried for cases diagnosed by a board-certified pathologist as usual interstitial pneumonia (UIP) in lung wedge biopsies performed for evaluation of diffuse parenchymal disease or therapeutic resections for lung cancer. Tissue termed “early UIP” was selected by a board-certified pulmonary pathologist to have relatively normal-appearing lung parenchyma with mildly thickened septa and small peripheral foci of scarring but not rising to the level of chronic interstitial pneumonia. When viewed in isolation, these features were insufficient for a diagnosis of UIP, but a definitive diagnosis of UIP was established based on findings elsewhere in these lungs. Normal lungs from deceased individuals rejected for lung transplantation were obtained from Gift of Life Michigan or from the Department of Pathology. Archived samples from lung explants with a diagnosis of UIP at Hannover Medical School were examined by transmission electron microscopy. The use of these samples was approved by the local ethics committee at Hannover Medical School or the University of Michigan Institutional Review Board.

### Immunostaining and Fluorescence In Situ Hybridization

Cross-sections of all mouse lungs and of human lung tissue measuring 1.5 cm in diameter were stained, imaged and analyzed in their entirety. Immunostained was performed as previously described(12). Briefly, tissue was deparaffinized, rehydrated, boiled in Target Retrieval Solution (Agilent S169984-2) or 1X sodium citrate buffer (10 mM, 0.05% Tween 20, pH 6.0), blocked with 5% donkey serum in 1X Tris-buffered saline with 0.05% Tween (TTBS), and incubated with primary antibodies overnight at 4°C. Primary antibodies included CCSP/SCGB1A1 (Seven Hills Bioreagents WRAB-3950), CD45 (R&D Systems MAB1430), CDKN2A/p16INK4a (Abcam ab108349), CLDN4 (Invitrogen 364800), GFP (Abcam ab13970), HOPX (Novus NBP1-92003), HTII-280 (Terrace Biotech TB-27AHT2-280), Ki67 (Invitrogen 50-5698-82), KRT5 (BioLegend 905504/095904 or Abcam ab321868), KRT8/TROMA-1 (Iowa Hybridoma AB531826), KRT17 (gift of Dr. Pierre Coulombe, University of Michigan or Abcam ab53707), LAMP3 (Synaptic Systems 391005), PDPN (Iowa DHSB 8.1.1), RAGE (R&D Systems AF1145), RFP (Rockland Immunochemicals 600-401-379 or Synaptic Systems 390004), SCGB3A2/UGRP1 (R&D Systems AF3545), or SFTPC/proSPC (Sigma-Aldrich AB3786). Secondary antibodies were applied for 1 h at room temperature. Secondary antibodies included Alexa Fluor 488-conjugated anti-guinea pig (Jackson ImmunoResearch 706-545-148), Alexa Fluor 647-conjugated anti-chicken (Jackson ImmunoResearch 703-605-155), Alexa Fluor 488-conjugated anti-rabbit (Jackson ImmunoResearch 711-545-152), Alexa Fluor 647-conjugated anti-goat (Jackson ImmunoResearch 705-605-147), Alexa Fluor 647-conjugated anti-rabbit (Jackson ImmunoResearch 711-605-152), Alexa Fluor 647-conjugated anti-syrian hamster (Jackson ImmunoResearch 107-605-142), Cy3-conjugated anti-chicken (Jackson ImmunoResearch 703-165-155), Cy3-conjugated anti-guinea pig (Jackson ImmunoResearch 706-165-148), Cy3-conjugated anti-rabbit (Jackson ImmunoResearch 711-166-152), Cy3-conjugated anti-rat (Jackson Immunoresearch 712-166-150), Cy5-conjugated anti-mouse (Jackson ImmunoResearch 715-175-150), Cy5-conjugated anti-rat (Jackson ImmunoResearch 712-175-150), FITC-conjugated anti-rat (Jackson ImmunoResearch 712-095-153), and FITC-conjugated anti-syrian hamster (Jackson ImmunoResearch 107-095-142). Sections were stained with DAPI (Thermo Fisher 62248) and mounted with Prolong Gold Antifade Reagent (Invitrogen P36930). Fluorescence *in situ* hybridization (FISH) was performed using RNAscope Multiplex Fluorescent Reagent Kit v2 (ACDBio 323110), *KRT17* probe (ACDBio 463661-C1), *tdTomato* probe (ACDBio 317041-C1), *Sftpc* probe (ACDBio 314101-C3), and *Sprr1a* probe (ACDBio 426871-C3) according to the manufacturer’s instructions, followed by immunostaining as previously described(80). Fluorescent images were acquired using a BX53 microscope (Olympus) at 20X. Focus points were placed across the tissue and multichannel tiles were acquired automatically and stitched into a single composite image. Exposure settings were set based on staining with secondary antibody alone and were constant between conditions but were typically less for insets. Hematoxylin, eosin, alcian blue, and trichrome staining were performed following standard protocols and imaged using Aperio AT2 or Aperio GT450 slides scanners (Leica Biosystems).

### Transmission Electron Microscopy

Samples were processed as previously described(97). Briefly, tissue was fixed by immersion with 0.1% glutaraldehyde, 4% paraformaldehyde in 0.2M HEPES buffer, post-fixed with 4% OsO4 and dehydrated in a series of increasing acetone concentration before embedding in Epoxy resin. Semithin sections (1 µm thickness) were cut and stained with toluide blue for initial light microscopic investigations. Tissue blocks with slight to moderate fibrotic remodeling (e.g. as characterized by thickened interalveolar septa and/or FF with otherwise maintained acinar microarchitecture and identifiable alveolar ductal and alveolar airspaces) were selected and ultrathin (80 nm) sections were cut and post-stained with lead citrate. Tissue was imaged by transmission electron microscopy (Morgagni 268) equipped with a 2×2K digital camera (Olympus Soft Imaging Solutions).

### Cell Culture

#### Primary Mouse AT2 Cells

AT2 cells were isolated from *S*ftpcCreERT2;mTmG or wildtype C57BL/6 mouse lungs as previously described(88). Briefly, lungs were harvested and digested using dispase (Corning 354235) and mechanical disruption and passed through 100 and 40 µm filters. Cells were incubated in FACS buffer (RPMI + 2% FBS) and stained with CD45 (BD Biosciences 553081), CD31 (BioLegend 102423), EPCAM (BioLegend 118210), and Lysotracker Deep Red (Invitrogen L12492) for 1 h and sorted using a Bigfoot Spectral Cell Sorter (Thermo Fisher). The AT2 fraction was defined as CD45-, CD31-, EPCAM+, and Lysotracker+. AT2 cells were seeded into serum-free, feeder-free alveolospheres in Matrigel (Corning 354234) and expanded in alveolar maintenance media(165). AT2 cell purity (>90%) was determined by immunostaining for proSPC (Sigma-Aldrich AB3786) immediately post-sorting and after each passage. AT2 cells, freshly isolated or after expanding, were cultured in 2D as previously described(166). Three independent experiments, each with 2-3 technical replicates, were performed.

#### Human iPSC-derived AT2 Cells

Human iPSC-derived (i)AT2 cells (clone SPC2-ST-B2), generated as described(167), were cultured as alveolospheres in Matrigel (Corning 354234) droplets covered with CKDCI medium [3 µM CHIR99021 (“C” ; Tocris 4432), 10 ng/mL rhKGF (“K”; R&D Systems 251KG010), 50 nM Dexamethasone (“D”; Sigma D4902), 0.1 mM 8BrcAMP (“C”; Sigma B7880), and 0.1 mM IBMX (“I”; Sigma I5879) in complete serum free differentiation base medium (“cSFDM”)] and passaged as described(^158^). After passaging, cells were cultured in CKDCI for 9-12 days, then treated with 5 ng/ml recombinant human TGFβ1 (PeproTech 100-21) in CKDCI or DCI media for 24 h. In indicated experiments, 10 mM SB431542 (Abcam ab120163), 10 mM MI-503 (MedChemExpress HY-16925), 10 µM A-485 (Cayman Chemical 24119), 10 µM SGC-CBP30 (Cayman Chemical 14469), or 10 mM SIS3 (MedChemExpress HY-13013) was added concurrently for 24 h. DMSO was used as a vehicle control. Cells were harvested for qPCR, RNA-seq, immunoblotting, or CUT&RUN, as described below. All iAT2 cell experiments were performed in at least 3 biological replicates from separate passages of a single iAT2 line.

#### Human Basal Cells

Human basal cells were isolated and cultured as previously described(168). Briefly, adult human trachea, obtained from Gift of Life Michigan, was resected and minced for 2 min into chunks <1mm^3^. Minced tissue was resuspended in Matrigel (Corning #356234), spotted into 20 µL droplets on a 24-well plate and cultured in Airway Organoid Media: DMEM/F12 (Corning 10092CV), 1X N2 Supplement (Gibco 17502048), 1X B-27 Supplement (Gibco 17504044), 0.05% Bovine Serum Albumin (Sigma-Aldrich A9647), 1X Penicillin-Streptomycin (Gibco 15140122), 100 ng/mL Ascorbic Acid (Sigma-Aldrich A4544), 0.4 µM Monothioglycerol (Sigma-Aldrich M1753), 500 ng/mL hFGF10 (R&D Systems 345FG), 1 µM A 83-01 (APExBIO A3133), 100 ng/mL hNoggin (R&D Systems 6057NG) and 10 µM Y-27632 (APExBIO 3008). Media was exchanged every 3-4 d. After 30 d, organoids were dissociated from Matrigel by mechanical dissociation (pipetting) and digested in TrypLE Express (Gibco 12605010) for 30 m, followed by quenching with equal volume DMEM/F12. Cells were pelleted by centrifugation (500g, 5 m) and resuspended at 200 cells/µl of Matrigel. Cells were passaged 3 times. To generate basal cell lines, established airway organoids were mechanically disrupted into small fragments and plated on tissue culture plastic containing a layer of mouse 3T3-J2 irradiated feeder cells (ATCC 30-2030). After 10 d of outgrowth, cells were treated with 0.25% Trypsin-EDTA (Gibco 25200056) and passaged at a ratio of 1:2 for the first passage and 1:20 for every subsequent passage. After 4 passages, RNA was harvested and qPCR was performed.

#### CRISPR-Cas9 Editing of KMT2A

A single guide (sg)RNA sequence targeting human *KMT2A* (caccGCAAAGGGTGGGATGCACAG) was designed using CRISPick (Broad Institute)(169). The *KMT2A* sgRNA or a non-targeting sgRNA (caccGTTCATTTCCAAGTCCGCTG) was cloned into plentiCRISPR v2 (Addgene 52961)(170). The cloned constructs were sequenced by Plasmidsaurus and used by the University of Michigan Vector Core to prepare lentiviral particles. Briefly, conditioned media from HEK293T cells transfected with the vector was concentrated and the lentiviral particles were resuspended in CKDCI medium with a titer of 10^6^-10^7^ TU/ml. iAT2 cells were dissociated into a single cell suspension as with passaging.. Single cells were incubated with the lentivirus (MOI=10) in CKDCI media at 37 °C, 5% CO_2_ for 4 h, pelleted, resuspended in 3D Matrigel, and cultured in CKDCI + 10 µM Y-27632 (Tocris 1254). 48 h after transduction, 1 mg/ml of puromycin (Gibco A1113803) was added to the media and Y-27632 was removed. After selection for 7 d, the alveolospheres were passaged and cells replated and cultured in CKDCI without puromycin. Of note, the line with the *KMT2A* sgRNA tended to grow more slowly. Genomic DNA was subjected to Sanger sequencing by Eurofins.

#### Quantitative Real-time (q)PCR and Bulk RNA Sequencing

To isolate total RNA from iAT2 cells, 1 mL of TRIzol (Invitrogen 15596018) was added to each well of a 24-well plate, and RNA extracted according to the manufacturer’s instructions. For mAT2s, 350 uL of 2-Mercaptoethanol (Sigma-Aldrich M3148) in Buffer RLT (Qiagen 74106) was added to each well and RNA extracted according to the manufacturer’s instructions. RNA was quantified using the Qubit RNA High Sensitivity Assay kit (Invitrogen Q32855) and Qubit 4 Fluorometer (Invitrogen Q33238). The QuantiTect Reverse Transcription Kit (Qiagen 205313) was used to make cDNA using 100-500 ng (iAT2) or 50-75 ng (mAT2) of total RNA. qPCR reactions were run using PowerUp SYBR Green Master Mix (Applied Biosystems A25742) on a QuantStudio 3 Real-Time PCR System (Thermo Fisher). The primer sequences used for qPCR reactions are listed in Table S7. Relative expression levels were calculated using the 2^−ΔΔCt^ method by normalization using the housekeeping genes *B2M* (*B2m)* or *Hprt* and were expressed as a “fold change” relative to the control sample. For bulk RNA-seq, RNA was submitted to the Advanced Genomics Core at the University of Michigan. RNA was assessed using Qubit RNA BroadRange Assay kit (Invitrogen Q10211) and TapeStation system (Agilent 5067-5576). Samples with RNA Integrity Numbers (RINs) of 8 or greater were processed using the NEBNext Poly(A) mRNA Magnetic Isolation Module (NEB E7490), NEBNext UltraExpress RNALibrary Prep Kit for Illumina (NEB E3330), and NEBNext Multiplex Oligos for Illumina Unique dual (NEB E6440). 170 ng of total RNA was converted to mRNA by polyA purification. The mRNA was then fragmented and transcribed into first strand cDNA using reverse transcriptase and random primers. The 3’ ends of the cDNA were adenylated and adapters were ligated. The products were purified and enriched by PCR to create the final cDNA library. Final libraries were checked for quantity and quality by Qubit dsDNA assay kit (Invitrogen Q33231) and DNA X-Mark Chip (Revvity Health Sciences 50-210-4506). The samples were pooled and sequenced on the Illumina NovaSeqX 10B Paired-end 150bp (Illumina 20085594) at a depth of 40 million reads per sample. BCL Convert (Illumina) was used to generate de-multiplexed Fastq files.

#### Immunoblotting

To prepare whole cell lysates, cells were homogenized in a lysis buffer containing 50 mM Tris (pH 7.4) (Fisher Scientific BP152), 150 mM NaCl (Fisher Scientific S640-3), 5 mM EDTA (Promega PRV4231), 0.5% NP-40 (Thermo Fisher 85142), 0.5% sodium deoxycholate (Sigma-Aldrich D6750), 1X protease inhibitor cocktail (Sigma-Aldrich P8340), and 1X phosphatase inhibitor cocktail (Sigma-Aldrich P5726). For the isolation of cytoplasmic and nuclear fractions, Nuclear Complex Co-IP kit (Active Motif 54001) was used. Cell lysate protein was quantified using a *DC* Protein Assay kit (Bio-Rad 5000112). Laemmli (Bio-Rad 1610747) + 2-Mercaptoethanol (Sigma-Aldrich M3148) was added to each sample and the samples heat denatured for 95°C for 10 min. Equal amounts of total protein were loaded onto a 4–15% Mini-PROTEAN TGX Precast Protein Gel (Bio-Rad 4561084) and transferred. Membranes were blocked for 2 h at room temperature in 5% milk, then primary antibodies were added overnight at 4°C on a rocker. After washing (5 min x 5), membranes were incubated with secondary antibodies for 1 h at room temperature with gentle rocking. Primary antibodies used include KRT17 (abcam ab53707), pSMAD2/3 (Cell Signaling 8828), pSMAD2/3 (Cell Signaling 3101), SMAD2/3 (abcam ab207447), β-actin (GenScript A00702), Histone H3 (Cell Signaling 9715), and KMT2A (Cell Signaling 14197). Secondary antibodies included anti-mouse HRP (Invitrogen 31430), anti-rabbit HRP (Invitrogen 31460), and anti-rat HRP (Invitrogen 31470). Images were acquired on an Amersham Imager 600 (GE) or a ChemiDoc Imaging system (Bio-Rad).

#### CUT&RUN

iAT2 cells cultured in 3D Matrigel in CKDCI media were treated with 5 ng/mL recombinant human TGFβ1 (PeproTeech 100-21) for 24 hr. Cells were trypsinized into a single cell suspension as per passaging. CUT&RUN and library prep were performed using CUTANA kits (EpiCypher 141048 and 141001) according to the manufacturer’s instructions. Antibodies used included H3K4me3 (EpiCypher 130060) and H3K27ac (EpiCypher 130059). The quantity and quality of the ensuing libraries were analyzed by Qubit (Invitrogen Q33231) and X-Mark Chip (Revvity Health Sciences 502104506). The samples were pooled and sequenced on the Illumina NovaSeqX 10B Paired-end 150bp (Illumina 20085594). BCL Convert (Illumina) was used to generate de-multiplexed Fastq files.

#### LDH Assay

iAT2 cells were seeded into 3D Matrigel and cultured in CKDCI + 10 µM Y27632 (Tocris) for 72 h. The cells were maintained in CKDCI (without Y-27632) for an additional 10-11 d, then treated with 5 ng/mL recombinant human TGFβ1 (PeproTech 100-21) in DCI medium for 48 h. An LDH assay (Abcam ab65393) was performed on conditioned media according to the manufacturer’s instructions. CKDCI or TDCI (i.e., TGFβ1 in DCI) medium was added at equal volume to empty wells (without cells) to serve as background controls for the assay. The reaction was allowed to incubate for 30 min at room temperature, then the plate was read at 450 nm using a SpectraMax M3 Microplate Reader (Molecular Devices) and analyzed with SoftMax Pro 7 software (Molecular Devices). The experiment was performed 3 times, each with 3 technical replicates.

#### Image Analysis

QuPath software (https://qupath.github.io/) was used to quantify the proportion of double positive cells and number of cells per µm^2^. Cells were detected using the DAPI channel at a cell expansion setting of 2 µm. To detect cells positive for an indicated marker, representative images were trained using the single measurement classifier tool and the classifier applied to all detected cells in the lung. Because *KRT5−/KRT17+* cells were rare, each KRT17+ cell detected was manually inspected to determine whether it was KRT5+. To quantify recombination efficiency of *S*ftpcCreERT2;mTmG mice, rectangular regions of interest consisting of approximately 1400 cells were drawn in each lobe and classifiers trained to detect proSPC+, GFP+, and proSPC+ GFP+ positive cells.

#### scRNA-seq Analyses

All publicly available scRNA-seq datasets were downloaded from the Gene Expression Omnibus (GEO) and processed in R using Seurat v5.3.1 (See Table S2). Where author-provided cell-type annotations were supplied as separate metadata files, these were matched to cells by barcode and added to the Seurat object. Author annotations were retained as the cell identities for all downstream analyses except for murine lung datasets, in which label transfer was performed. Unless otherwise specified, raw count matrices were log-normalized, 2,000 highly variable features were identified by variance-stabilizing transformation, expression values were scaled, and dimensionality reduction was performed by principal component analysis (PCA) followed by UMAP or t-SNE projection. Cluster identities were either extracted directly from the published annotations or manually assigned based on marker gene expression, as specified in the figure legend. Processed objects were deployed as interactive ShinyCell2 applications, and all embedding, dot-plot, and feature-plot panels shown were exported from these applications. Dot plots display mean scaled expression and the proportion of cells with detectable expression within each annotated identity. Feature plots display log-normalized expressions. For all heatmaps and dot plots shown, gene expression was scaled across rows.

Modules for TGFβ, senescence, and proteostasis were generated from published gene sets(88). To create the human IPF *KRT5−/KRT17+* gene module, the top 50 differentially expressed genes in *KRT5−/KRT17+* cells compared to all lung epithelial cells was obtained from GSE135893(64). Before scoring, module genes were intersected with the mouse genome such that symbols without a corresponding mouse gene were dropped (and not counted as zero). Likewise, a module of the top 50 basal cell genes (ranked by fold change and with p-adj < 0.05) expressed by IPF *KRT5−/KRT17+* cells was created from GSE135893(64). To create the human IPF *KRT5−/KRT17+* Discriminative vs. Mouse module, the top 50 genes upregulated in *KRT5−/KRT17+* cells relative to AT2 cells (ranked by log_2_FC > 0) was obtained from GSE135893(64) and converted to mouse symbols. Genes upregulated in cluster 1 (log_2_FC > 0 and p-adj < 0.05) relative to all other cells in a murine pneumonectomy dataset (GSE138585 and GSM2858341)^82^ were then subtracted from the list of top 50 genes – where cluster 1 refers to the authors’ original clustering (clusters_8) – yielding 32 *KRT5−/KRT17+* discriminative marker genes that murine transitional cells do not generally express. For the mouse lung transitional state module, the top 50 DEGs ranked by fold change (all p<0.05) common to the bleomycin and LPS datasets were used(63, 80). For the kidney, skin, and intestine datasets, genes differentially expressed between the injury-emergent cell type of interest and all other epithelial cell types from that tissue were identified with Seurat’s FindMarkers (using a Wilcoxon rank-sum test and a minimum detection threshold of 25% of cells) and compiled into a module. The modules were then scored on the epithelial cell types of each lung dataset using AddModuleScore. For all modules, genes with an adjusted p-value (padj) < 0.05 were ranked by log fold change (log_2_FC > 1) and per-cell scores were computed with AddModuleScore and are displayed on the corresponding embedding in arbitrary units (A.U.). Module scores were summarized per cell type and displayed as dot plots generated with ggplot2. Scale Bars indicate Cell State or Pathway Module scores (in A.U.). Cell State Module scores > 0 reflect expression of module genes above a control-gene background, which AddModuleScore subtracts to account for differences in overall expression level. Dot plots were generated through ShinyCell2 applications. For plots of marker gene expression by identified cell types, scale bars indicate the average gene expression as a z-score. See Tables S3, S4, and S6 for gene lists for each module.

Global transcriptional similarity was quantified with scTOP(100). Unless otherwise specified, count matrices were exported from the Seurat objects and subset to the cell types being compared as MatrixMarket files with accompanying gene, cell, and metadata tables. Reference bases were built with create_basis(). To confirm that each basis resolved its own cell identities, the reference dataset was projected back onto its own basis as a query. Query cells were then processed with process() and projected onto each basis with score(). Mean scTOP scores per cell type were displayed as heatmaps. Mouse genes were converted to one-to-one human orthologs with biomaRt. Bases were built from raw counts with *create_basis*, excluding cell types with fewer than 30 cells. Human AT1, AT2, Transitional AT2, *KRT5−/KRT17+*, Basal, SCGB3A2+, and SCGB3A2+ SCGB1A1+ cells from GSE135893^64^ were normalized with process and projected onto each basis. Projection scores were averaged per human lung cell type and displayed as heatmaps on a shared color scale.

A prior scRNA-seq meta-analysis(88) was re-analyzed. Cells were annotated using the published cluster assignments. From the eight-cluster annotation (clusters_8), clusters containing very few cells, all derived from a single study were removed. The remaining five clusters were renumbered 1–5. From the seven-cluster annotation (clusters_7), clusters were removed using the same criteria and the remaining clusters were renumbered 1–4. UMAP coordinates were computed on the full dataset before filtering and retained such that filtered cells occupy the same positions as in the complete embedding. The Bleomycin: *Sftpc* Lineage Trace dataset was obtained from GSE252588^44^. The AT2-tracer (GSM8003560) experiment was analyzed. Processing was performed using the code deposited by the authors on Zenodo. PCA and UMAP were computed, nearest-neighbor graphs constructed, and Leiden clustering applied at a resolution of 0.2. Contaminating immune clusters were removed before PCA and UMAP were recomputed. The Pneumonectomy: *Sftpc* Lineage Trace dataset was obtained from GSE138585 and GSM2858341^82^. Sample N0 cells from the dataset were derived from untreated knockout mice and were excluded as they were not relevant to the injury-associated states studied here. Repetitive Bleomycin: *Sftpc* Lineage Trace dataset was obtained from GSE243252(99). In Seurat, non-epithelial cells were identified by low *Epcam* expression and removed together with cells expressing *Ptprc*, *Col1a1*, *Pecam1*, *Ms4a1*, *Mki67*, *Krt5,* and *Sox8* as published by the authors. PCA, Harmony correction, neighbor-graph construction, clustering (resolution 0.54), and UMAP were recomputed on the retained epithelial cells. The epithelial object was then re-clustered at a resolution of 0.4. Ciliated cells and doublets were removed. Clustering (at resolution 0.4) and UMAP were recomputed on the remaining cells. The resulting object was split by sample (orig.ident). For each sample, two variants were generated: one retaining the coordinates and cluster assignments carried over from the merged object and one re-embedded independently (log-normalization, variable feature selection, scaling, PCA, UMAP, and nearest-neighbor graph on the first 30 principal components) and clustered across resolutions 0.01–0.3. Module scores were added to all variants. The projection was clustered at a resolution of 0.08. For each of the above murine lung *Sftpc* lineage tracing datasets, cell clusters were assigned the scANVI class with the highest predicted probability. Predicted labels were visualized on a UMAP computed from the scANVI latent space and used for scTOP analysis.

#### Epigenetic Modifiers

To obtain a comprehensive list of epigenetic modifiers, we searched epifactors.autosome.org/genes for those that had the “function” of factors that write or erase histones or modify DNA, with the “modification” being (de)acetylation or (de)methylation, leaving 141 candidate factors. These factors, selected for a padj < 0.05, were ranked in order of fold change expression between *KRT5−/KRT17+* cells and all other epithelial cells(64).

#### Bulk RNA-Seq Analysis

Raw sequencing data were processed with the nf-core/rnaseq pipeline (v3.21.0; Nextflow 24.10.5). Adapter and quality trimming were performed with Trim Galore (v0.6.10; Cutadapt v4.9), and read quality was assessed with FastQC (v0.12.1). Reads were aligned to the human reference genome (GRCh38, Ensembl release 115; soft-masked primary assembly) with STAR (v2.7.11b), and gene-level quantification was performed with RSEM (v1.3.1). Alignment and quantification quality were evaluated with RSeQC (v5.0.2), Qualimap (v2.3), Picard (v3.1.1), dupRadar (v1.32.0), and MultiQC. Differential gene expression analysis was performed in R (v4.5.1) with DESeq2 (v1.48.2). RSEM expected counts were rounded to integers, and genes with counts ≥10 in at least three samples were retained. The model design was ∼condition, and pairwise contrasts (TDCI vs. CKDCI, and TDCI+MI-503 vs. TDCI) were extracted with independent filtering enabled. For each contrast, a positive log_2_FC indicates higher expression in the first-named group. Genes with Benjamini–Hochberg padj < 0.05 were considered differentially expressed.

#### CUT&RUN Analysis

Raw sequencing reads were processed with the nf-core/cutandrun pipeline (v3.2.2)(171) executed under Nextflow (v25.04.8) with Singularity containers. Read quality was assessed with FastQC (v0.12.1), and adapters and low-quality bases were trimmed with Trim Galore (v0.6.6; cutadapt v2.10) using two-color chemistry quality trimming (Phred ≥ 20). Trimmed reads were aligned to the GRCh38 human reference genome with Bowtie2 (v2.4.4) using pipeline default parameters. Alignments were sorted, indexed, and filtered with SAMtools (v1.17), and duplicates were marked with Picard MarkDuplicates (v3.1.0) and removed from IgG control samples only, per pipeline defaults. Reads overlapping blacklisted regions were excluded using BEDTools (v2.30.0), using the GRCh38 blacklist distributed with the nf-core/cutandrun pipeline (assets/blacklists/GRCh38-blacklist.bed) and as --normalisation_mode None. Peak calling was performed with MACS2 (v2.2.7.1) and SEACR (v1.3) using matched IgG samples as background controls. For H3K4me3, MACS2 narrow peaks were used. For H3K27ac, SEACR Stringent peaks were used for downstream analyses. For SMAD2/3, MACS2 narrow peaks were used. Differential binding analysis was performed using DiffBind (v3.6). Read counts within consensus peaks were normalized using the trimmed mean of M-values (TMM) method, and differential binding between conditions (TDCI vs CKDCI) was assessed with DESeq2 as implemented in DiffBind, using a design accounting for experimental run (∼Replicate + Condition). For SMAD2/3, TMM normalization was computed on reads in large genomic background to avoid bias from global shifts in SMAD2/3 occupancy between conditions. Differentially bound regions were annotated with ChIPseeker (v1.42.0) using the UCSC hg38 known-gene models (TxDb.Hsapiens.UCSC.hg38.knownGene), with gene symbols mapped via org.Hs.eg.db. Each peak was assigned a single genomic feature; where a peak overlapped multiple features, one was selected according to ChIPseeker’s default priority order: promoter (± 3 kb of the TSS), 5′ UTR, 3′ UTR, exon, intron, downstream, and distal intergenic. Coverage tracks were normalized using the same scaling factors applied in the differential binding analysis, so that browser tracks, metagene profiles, and heatmaps are quantitatively consistent with the statistical results. Browser tracks images were created using Integrative Genomics Viewer software.

To determine enrichment of H3K4me3 in AT2 cells, we retained MACS2 peaks called in at least two of three CKDCI replicates (17,916 peaks). A gene was considered marked if a peak overlapped ±2 kb of any of its annotated transcription start sites (TxDb.Hsapiens.UCSC.hg38.knownGene). The proportion of marked genes among the 189 genes upregulated in *KRT5−/KRT17+* cells^64^ (adjusted p < 0.05, lnFC > 0.58) was compared with that of all other protein-coding genes using a two-sided Fisher’s exact test. The same analysis was performed for H3K27ac (SEACR stringent peaks) and SMAD2/3 (MACS2 narrow peaks). Duplicate-marked target alignments were converted to genome-wide coverage tracks using deepTools (v3.5.6) bamCoverage with the inverse of the DiffBind-derived TMM normalization factors supplied as per-sample scaling factors (-- scaleFactor, --normalizeUsing None, --binSize 50). No additional read-depth or library-size normalization was applied, so that track amplitudes remain on the same quantitative scale used for differential testing. Scaling factors for control replicates 1–3 and TGF-β replicates 1–3 were 0.5382, 1.0558, 1.4723 and 0.7292, 1.6281, 3.0525 for H3K4me3, and 0.9075, 0.6270, 0.9781 and 1.2223, 1.3837, 1.4312 for H3K27ac, and 1.0755, 0.9243 and 0.9992, 1.0127 for SMAD2/3 (control and TGF-β replicates 1–2). Scaled replicate tracks were averaged within each condition using bigwigAverage (50 bp bins). Signal was quantified with computeMatrix reference-point (reference point = TSS, 3 kb upstream to 3 kb downstream, 50 bp bins, --skipZeros) over TSSs from the pipeline reference annotation, retaining 87,053 (H3K4me3) and 87,050 (H3K27ac) of 95,582 regions with non-zero signal. Heatmaps were rendered with plotHeatmap using the RdYlBu_r (H3K4me3) or RdBu_r (H3K27ac) color map. For each mark, both conditions were quantified within a single matrix and plotted on a common, automatically scaled color range, so signal intensities are directly comparable between conditions. The upper panel shows the mean normalized coverage profile across all retained regions for each condition (arbitrary units); the lower heatmaps show per-region signal, ordered by mean signal across the window. For H3K27ac, metagene profiles spanning the full gene body (TSS to TES, 5 kb scaled region body, ±3 kb flanks) were also generated using computeMatrix scale-regions with otherwise identical parameters. We defined a gene as having increased SMAD3 occupancy if any peak assigned to it showed a fold change >1.5 and FDR <0.05 (TGF-β vs. control; IgG-subtracted, spike-in–normalized DiffBind); genes without a SMAD2/3 peak counted as not increased. We compared the proportion of *KRT5−/KRT17+* genes with that of unchanged genes (|log2 fold change| <0.15, *P* >0.5; TGF-β vs. control, DESeq2), limiting both groups to genes expressed in iAT2 cells (baseMean ≥10), by Fisher’s exact test. To control for expression level, we repeated the test 200 times against unchanged genes sampled 5:1 within deciles of baseMean and report the median *P* value.

SMAD2/3 consensus peaks within 50 kb of a KRT5⁻/KRT17⁺ gene TSS (nearest gene, ChIPseeker; n = 330) were centered on their summits and split into promoter (TSS ±3 kb; n = 209) and distal (3–50 kb; n = 121) groups. Replicate-averaged TGF-β tracks (background-TMM SMAD2/3, TMM H3K4me3 and H3K27ac, IgG) were quantified with computeMatrix reference-point (±5 kb, 100 bp bins). Rows were sorted by mean SMAD2/3 signal within ±500 bp of the summit, and heatmaps were plotted with a separate color scale per mark (98th percentile; H3K4me3 capped at 700).

KRT5_KRT17_vs_allepi.csv and krt17_vs_at2.csv Supplemental Tables were downloaded(64). *KRT5−/KRT17+* marker genes were defined as *KRT5−/KRT17+* versus all epithelial cells fold change ≥ 1.5, padj<0.05. AT2 genes were defined as *KRT5−/KRT17+* versus AT2 fold change ≥ 1.5, padj<0.05.

#### Statistics

Analysis was performed using GraphPad Prism v10.4.1. Data are presented as mean ± SEM. p<0.05 was considered statistically significant. For qPCR and RNA-seq-based comparisons of gene expression between two conditions, we used a one-sample t-test (for comparisons normalized to a control set at 1) or a ratio paired t-test (for paired treatment/no-treatment comparisons) on log_2_fold change values, with the Benjamini, Krieger, and Yekutieli two-stage method used to control the false discovery rate across multiple comparisons within an experiment. Comparisons of expression between two independent, non-paired groups (e.g., basal cells versus TGFβ-treated iAT2 cells) were assessed by Welch’s t-test with the same multiple-comparison correction. Significance is denoted in the figures using asterisks (*q < 0.05, **q < 0.01, ***q < 0.001, ****q < 0.0001). Gene-level q-values for each comparison are provided in Table S5. Enrichment of marker gene sets (e.g., *KRT5−/KRT17+* or AT2 identity genes) among genes gaining or losing a given histone mark, or among genes inhibited or restored by a pharmacological treatment, was assessed by Fisher’s exact test, with odds ratios (OR) and p-values reported in the text or figure legend. The relationship between the magnitude of a chromatin mark change (e.g., H3K4me3 gain, divided into tertiles) and the magnitude of a downstream pharmacological effect was assessed by the Kruskal-Wallis test. Genome-wide comparisons of normalized CUT&RUN read counts (log_2_) between conditions, either at the level of individual marker genes or across gene sets, were assessed by the Mann-Whitney test, comparing the median change in signal for the gene set of interest to that of all other expressed/annotated genes.

## DATA

The bulk RNA-seq and CUT&RUN data from this study have been deposited at GEO using accession numbers GSE353612 and GSE353611, respectively. This paper analyzes existing scRNA-seq datasets, whose accession numbers are listed in Table S2.

## CODE

All code has been deposited online at https://github.com/ZemansLab and will be publicly available at the time of publication in a peer-reviewed journal.

## USE OF GENERATIVE AI

During the preparation of this work, the author(s) used Claude to perform literature searches and edit text for clarity and typographical errors. In addition, Claude was used for debugging, interpreting and editing scripts, and for writing codes and functions for scRNA-seq analyses. Afterwards, the authors reviewed and edited the manuscript and are responsible for all content.

## AUTHOR CONTRIBUTIONS

R.L.Z. conceived and administered the project. A.B. curated CUT&RUN, bulk RNA-seq, and scRNA-seq data. J.L., X.Z., R.G., A.B., Q.X., K.K., L.K., R.T., and R.L.Z. performed data analysis. J.L., X.Z., R.G., K.F., Q.S., C.T., H.P., J.F. C.W., and L.K. performed experiments. L.A. and D.N.K. supplied iAT2s and established protocols for its culture. C.O. and C-Y.L. provided input on CUT&RUN protocols, experimental design, and analysis. R.P.D. and N.F. performed and advised on the design of hyperoxia experiments. K.K. provided and interpreted normal and IPF lung samples. J.R.S. and T.F. provided primary human basal cells. J.L., X.Z., A.B., Q.S., and L.K. independently validated key results. J.L., X.Z., A.B., R.G., and R.L.Z. wrote the manuscript. J.L., X.Z., R.G., K.K., L.K., L.A., C-Y.L., D.N.K., and R.L.Z. edited the manuscript. All authors reviewed and approved the paper.

## ACKNOWLEDGEMENTS

We acknowledge support from the University of Michigan’s Bioinformatics, Advanced Genomics, Viral Vector, Histology (Dental School), and Digital Pathology Cores. We thank Linda Samuelson, Jeffrey Myers, Tushar Desai, Timothy Blackwell, Brigitte Gomperts, and Richard Simon for thoughtful discussions and Katherine Gallagher, Amrita Joshi, Dylan Cable, Xintong Zhai, Madeline Lyu, Nicholas Juul, Yue Zhao, and Mashrafi Bhuiyan for technical assistance. This study was supported by the following grants: NIH F31HL175884 (JL), R35HL160770 (RLZ), R01ES035710 (RLZ), T32HL007749 (RG), F32HL172379 (CT), R01NS134942 (C-YL), R01HL095993, U01HL148692, P01HL170952, U01HL175451, and U01HL152976 (DNK). iAT2 cell sharing was supported by N01:75N92025D00035 to DNK.

---

**Supplemental Figure 1.**
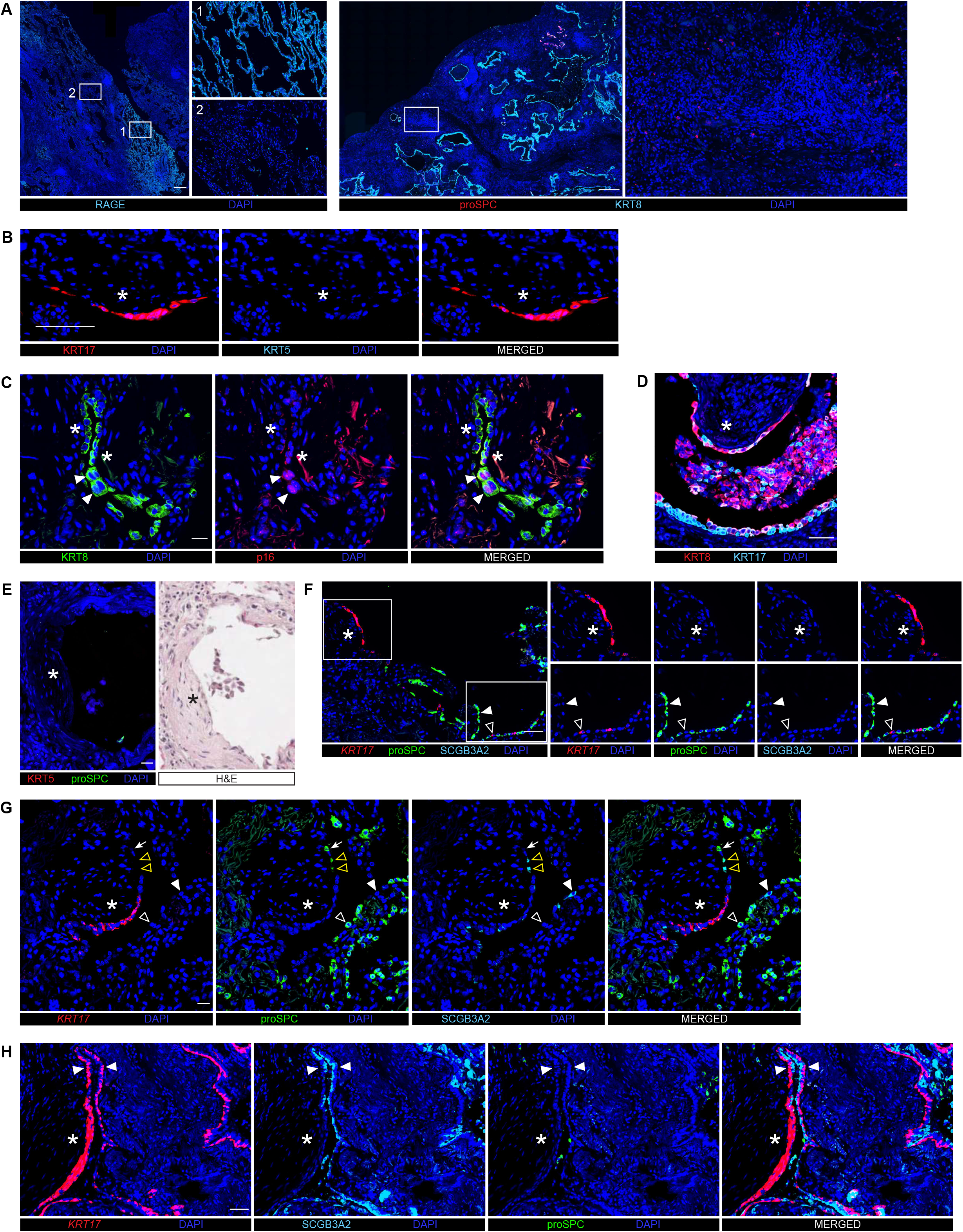
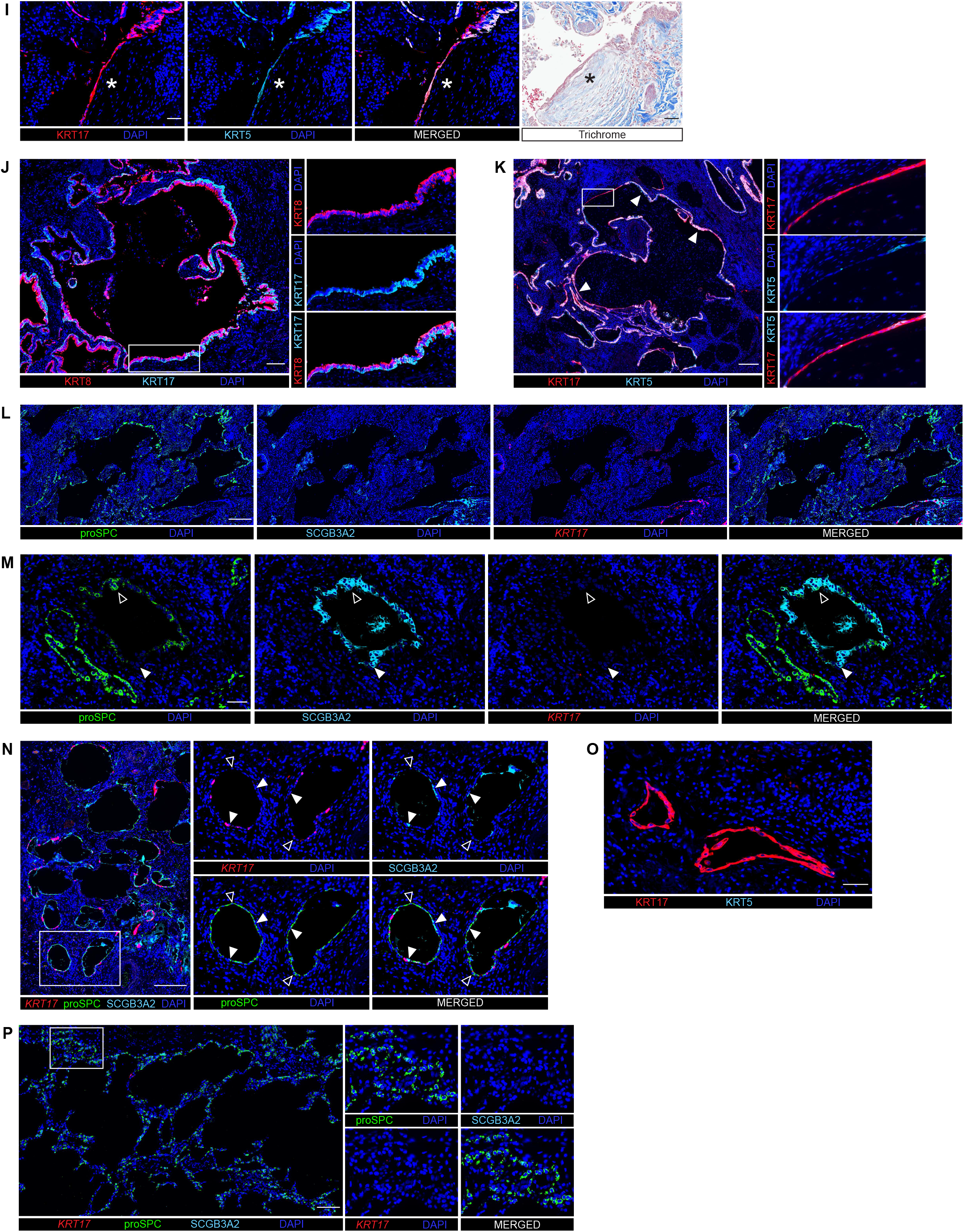
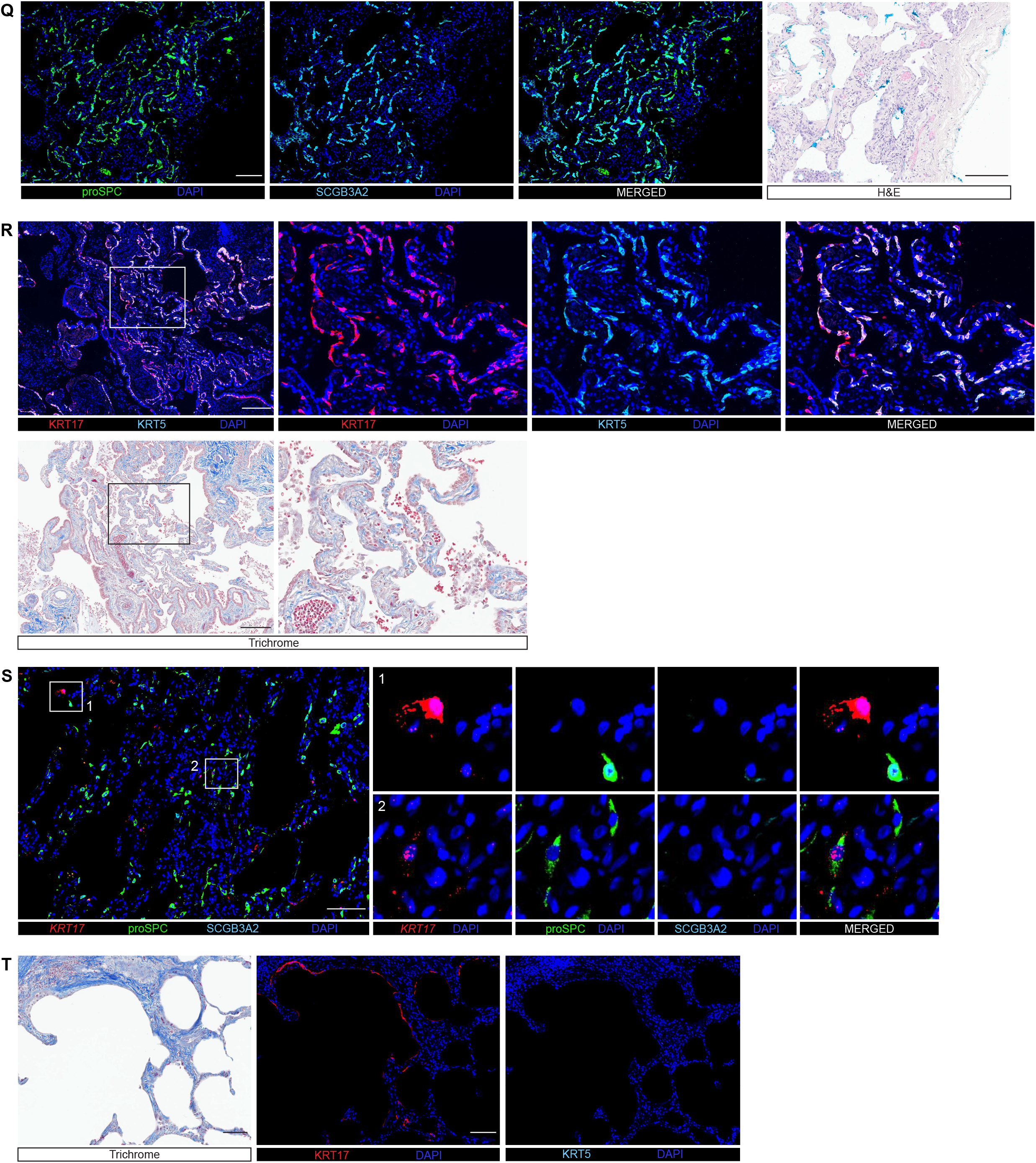
IPF lung is characterized by a microanatomically and molecularly haphazard epithelium in a severely remodeled environment. Human IPF lung stained with H&E, trichrome, or indicated antibodies and FISH probes. FF (*). (F) proSPC+ SCGB3A2-monolayer (closed arrowhead), *KRT17+* cells (open arrowhead). (G) proSPC+ SCGB3A2+ (open yellow arrowheads) and proSPC+ SCGB3A2-(white arrow) cells on FF. proSPC+ SCGB3A2-(open white arrowhead) and proSPC-SCGB3A2+ (closed white arrowhead) cells on adjacent epithelium. (H) Pseudostratified airways (arrowheads). (K) KRT17+ KRT5-monolayer (inset), pseudostratified epithelium (closed arrowheads). (M) proSPC+ SCGB3A2+ (open arrowhead), proSPC-SCGB3A2+ (closed arrowhead) cells. (N) proSPC+ SCGB3A2+ (closed arrowheads), proSPC+ SCGB3A2-(open arrowheads) cells. Scale Bars: 500 µm (A), 100 µm (B,J,P-R,T), 20 µm (C,E,G), 50 µm (D,F,H,I,M,O), 200 µm (K,L,N,S), and 4 mm (T). n≥3 patients.

**Supplemental Figure 2.**
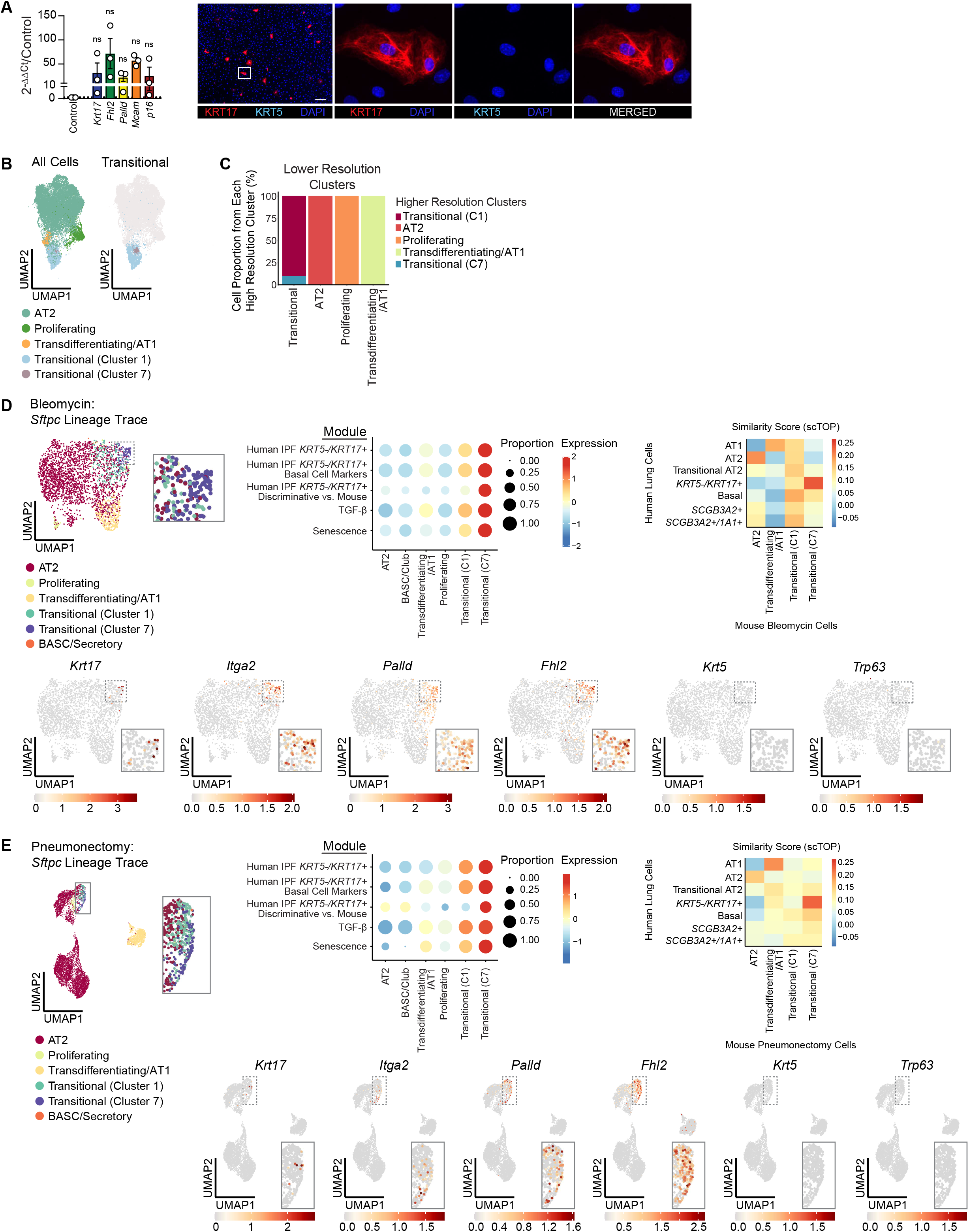
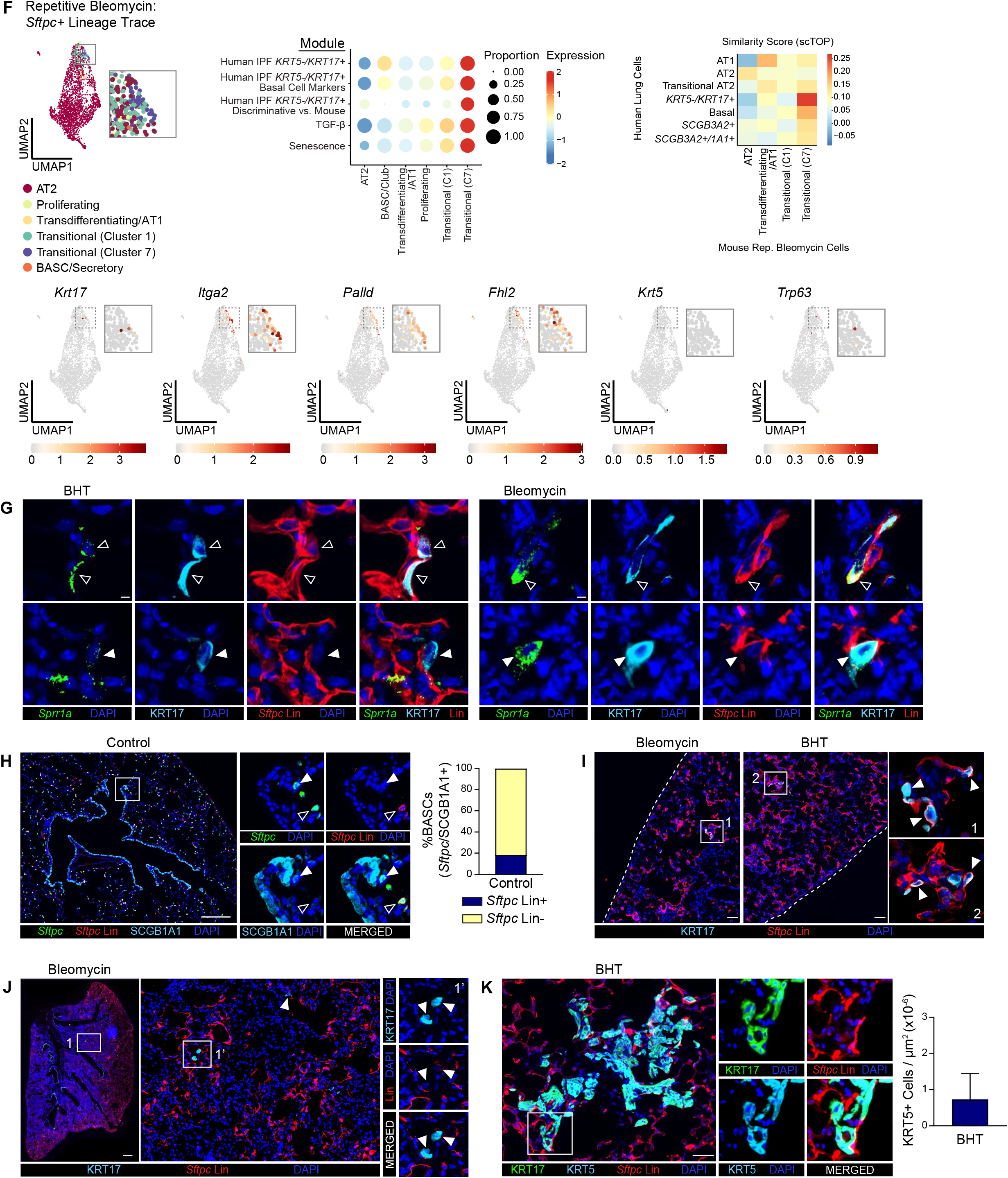
K*R*T5*−/KRT17+* cells arise from AT2 Cells. (A) Primary murine AT2 cells were cultured in 2D for 3d. qPCR (left panel) and immunostaining for indicated markers (right panel) was performed. qPCR: one sample ratio t-test (log_2_FC) with Benjamini, Krieger and Yekutieli correction. n=3. (B) UMAP of integrated scRNA-seq datasets(63, 80, 81), reproduced from prior publication(88) under a CC BY 4.0 license. (C) Reclustering of the integrated dataset in (B) at reduced resolution. Stacked bars show the proportion of cells from each high-resolution cluster assigned to each low-resolution cluster (x axis). (D-F) Analysis of *Sftpc* lineage-traced cells from the (D) bleomycin(44), (E) pneumonectomy(82), and (F) repetitive bleomycin(99) models. Label transfer from the integrated dataset (B) onto each dataset; UMAPs show the resulting cluster assignments. Dot plots show module scores for human *KRT5−/KRT17+* markers, basal cell markers expressed by *KRT5−/KRT17+* cells, human *KRT5−/KRT17+* versus mouse transitional discriminative genes, and TGFβ and senescence pathway genes. Heatmaps show mean scTOP similarity scores between each human lung epithelial cell type(64) and reference mouse lung cell types. UMAPs show expression of selected *KRT5−/KRT17+* and basal cell markers. (G-K) *S*ftpcCreERT2;mTmG mice were subjected to the indicated treatments and stained with the indicated antibodies and FISH probes. (G) *Sftpc* lin+ (open arrowheads) and lin-(closed arrowheads) *Sprr1a+* KRT17+ cells. (H) *Sftpc* lin-*Sftpc+* SCGB1A1+ cell (closed arrowhead) and *Sftpc* lin+ *Sftpc+* SCGB1A1-cell (open arrowhead). Percent of *Sftpc+* SCGB1A1+ cells that are *Sftpc* lin+ or lin-. (I) *Sftpc* lin+ KRT17+ cells (arrowheads). Dotted lines indicate pleura. (J) *Sftpc* lin-KRT17+ cells (arrowheads). (K) Number of KRT5+ cells per um^2^. Scale Bars: 5 µm (G), 200 µm (H), 50 µm (I,K), 500 µm (J). n=3-4 mice/group. Error bars represent Mean ± SEM.

**Supplemental Figure 3.**
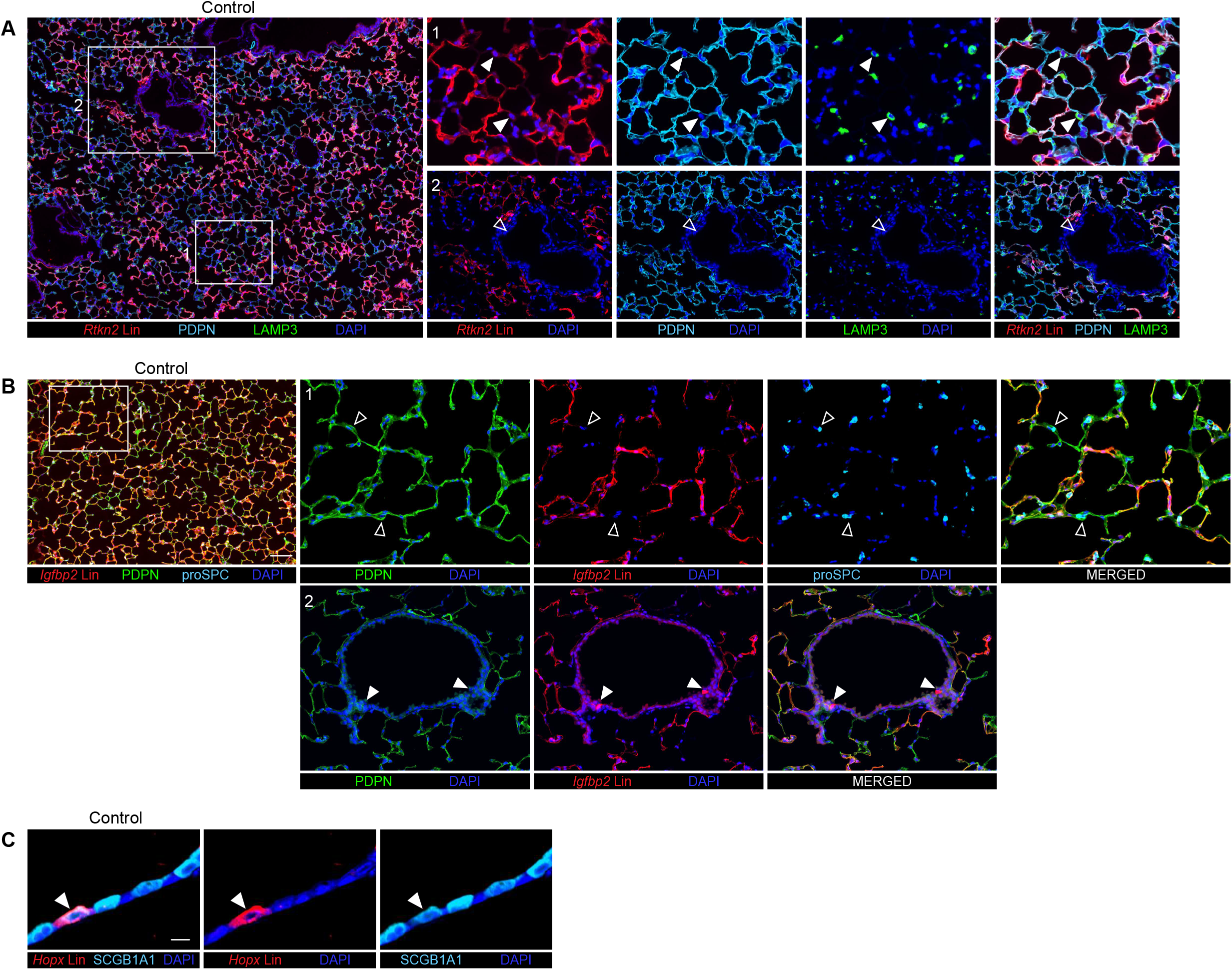
Specificity of AT1 Cre drivers for lineage tracing. (A-B) *Rtkn2CreERT2^-^;tdTomato* (A) and *Igfbp2CreERT2;mTmG* (B) mice were treated with tamoxifen and their lungs stained with the indicated antibodies. (A) *Rtkn2* lin-LAMP3+ cells (closed arrowheads) and *Rtkn2* lin-cells (open arrowhead). (B) *Igfbp2* lin-proSPC+ cells (open arrowheads) and *Igfbp2* lin+ cells (closed arrowheads). (D) *HopxCreER;mTmG* mice were administered tamoxifen and lungs immunostained for the indicated antibodies. *Hopx* lin+ SCGB1A1+ cell (arrowhead). Scale Bars: 100 µm (A,B) and 10 µm (C). n=2 mice/group.

**Supplemental Figure 4.**
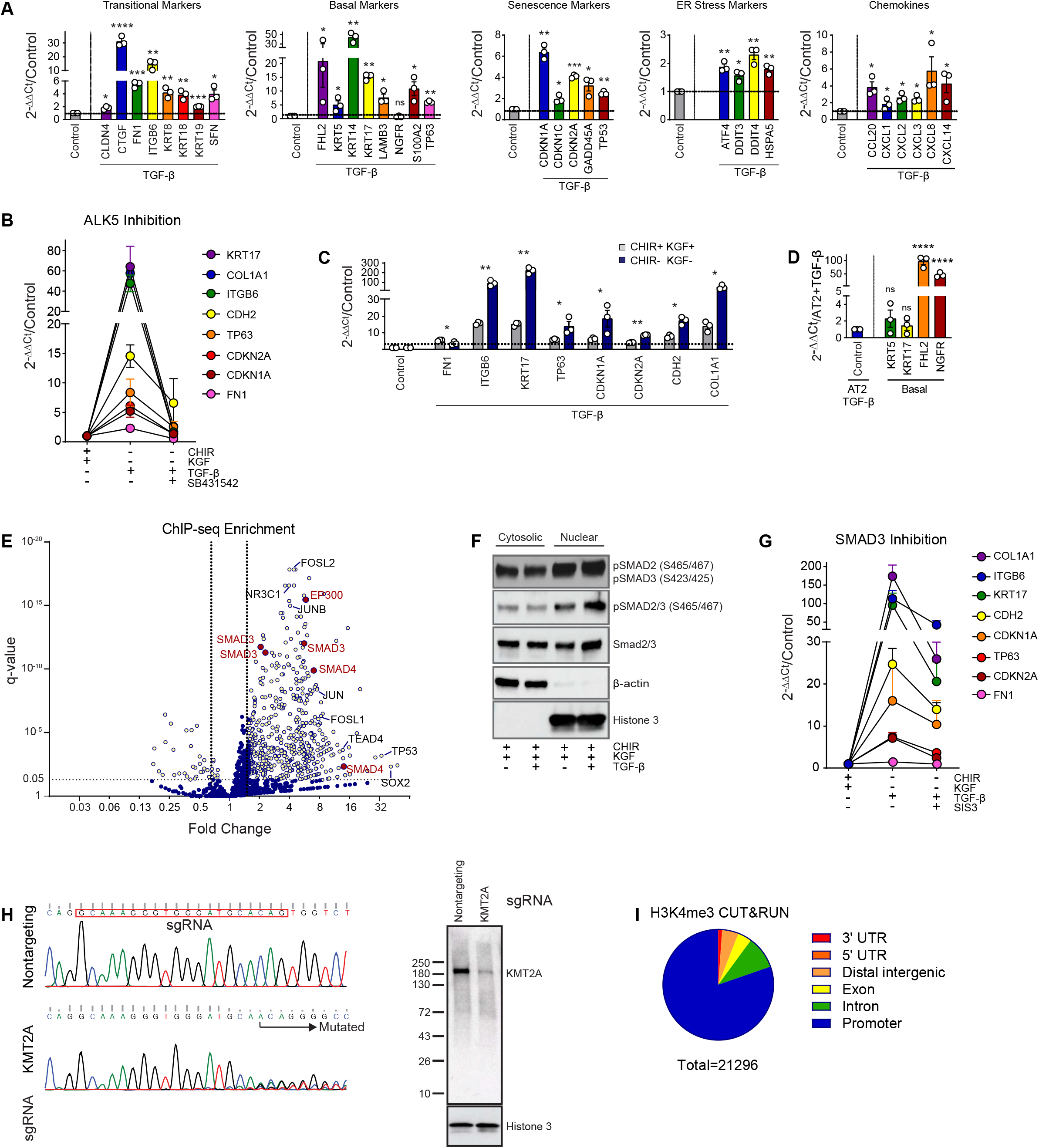
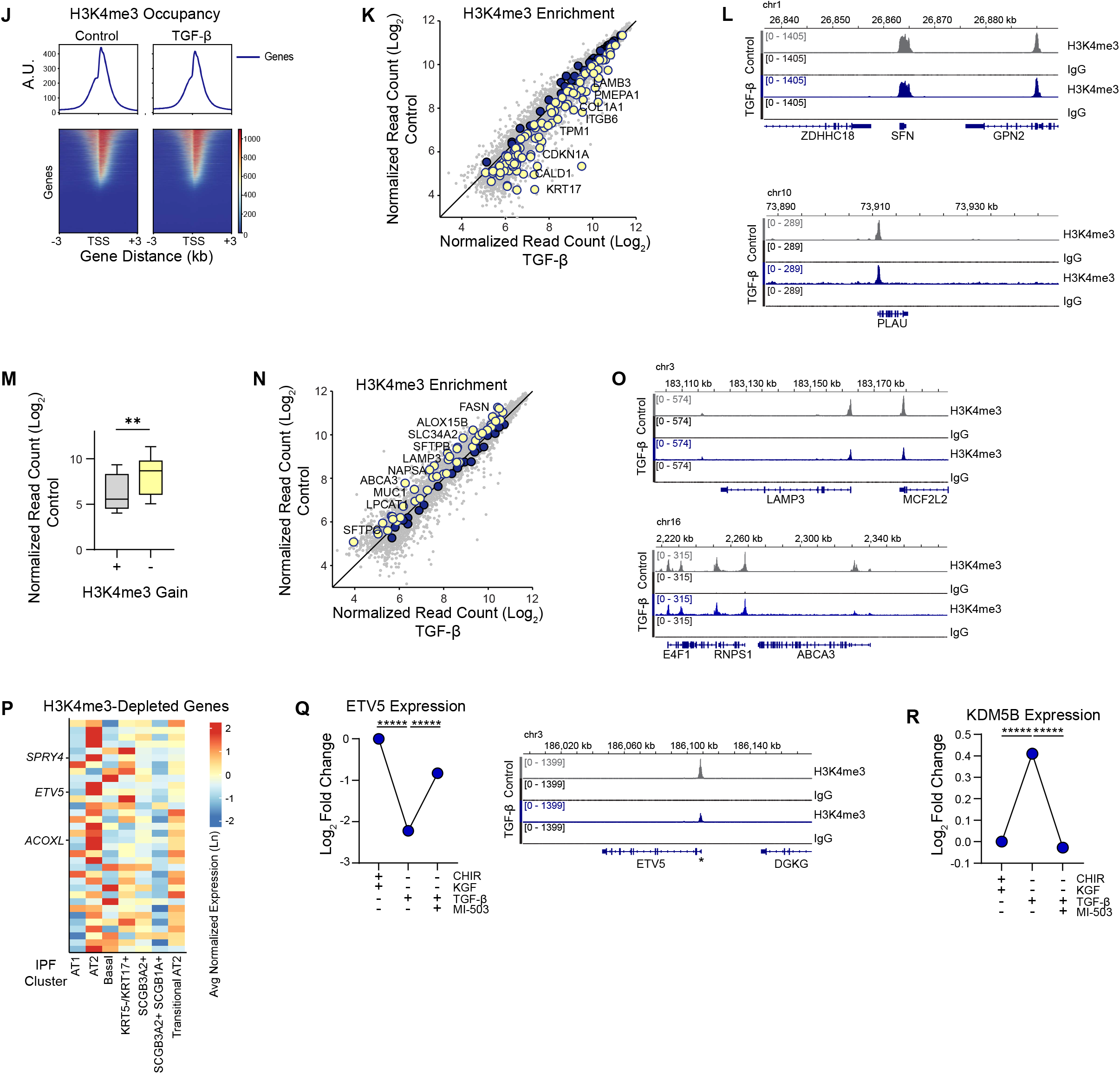
TGF-β/SMAD3 induces the *KRT5−/KRT17+* state via KMT2A-mediated H3K4 trimethylation. (A-D,G) Expression of indicated genes, as determined by qPCR, in iAT2 cells treated with TGF-β in the presence or absence of indicated inhibitors (A-C,G) or unstimulated basal cells compared to iAT2 cells treated with TGF-β (D). (E) Transcription factors and chromatin regulators enriched at *KRT5−/KRT17+* marker gene loci, as determined by a query of *KRT5−/KRT17+* marker genes against a database of publicly available lung epithelial ChIP-seq datasets(172). Red dots: factors of interest. Dotted lines: q=0.05, FC = 0.67,1.5. (F) Western blot of cytosolic or nuclear lysates of iAT2 cells treated with TGF-β. Representative of n=3 independent experiments. (H) iAT2 cells transduced with non-targeting or *KMT2A*-targeting lentiviral CRISPR-Cas9 and analyzed by Sanger sequencing of the targeted locus (left) and western blot (right). (I-Q) AT2 cells were treated with TGF-β and subjected to CUT&RUN with an H3K4me3 antibody or IgG. (I) Genomic distribution of all called H3K4me3 peaks across both control and TGF-β conditions, average of n=3. (J) Genome-wide H3K4me3 occupancy shown as TSS-centered profile plots and heatmaps (±3 kb) comparing untreated AT2 cells (Control) and TGF-β-treated (*KRT5−/KRT17+)* iAT2 cells. (K) Normalized H3K4me3 CUT&RUN read counts (log_2_) at all annotated peaks. Large dots: *KRT5-/KRT17* marker genes(64) with increased (yellow dots) or decreased (blue dots) H3K4me3 following TGF-β-treatment. No fold-change or p-value cutoff applied. *KRT5−/KRT17+* marker genes showed gene-set-level H3K4me3 gain following TGF-β compared to other genes (median log_2_FC 0.27 vs. 0.01; p = 1.6×10^-13^, Mann-Whitney test). 87.3% of *KRT5−/KRT17+* genes were marked by H3K4me3 in AT2 cells (OR 3.01 relative to other protein-coding genes, p=1.6×10^-8^ by Fisher’s Exact Test). (L,O,Q) Representative genome browser tracks showing mean H3K4me3 and IgG CUT&RUN signal at the indicated loci in untreated (Control) and TGF-β-treated (TGF-β) iAT2 cells. (M) *KRT5−/KRT17+* genes upregulated by TGF-β (FC>1.5, padj<0.05) and inhibited by MI-503 (FC<0.67, padj<0.05 versus TGF-β alone), stratified by H3K4me3 change: H3K4me3 gain (log_2_FC>0.20) versus no (log_2_FC<0.20). Box plots show baseline H3K4me3 signal in untreated AT2 cells. Box: interquartile range; line: median; whiskers: min-max. **p<0.01 by t-test. (N) Normalized H3K4me3 CUT&RUN read counts (log_2_) at all annotated peaks. Large dots: AT2 markers compared to *KRT5−/KRT17+* state(64), with decreased (yellow dots) or increased (blue dots) H3K4me3 following TGF-β-treatment. No fold-change or p-value cutoff applied. AT2 marker genes showed gene-set-level loss of H3K4me3 following TGF-β compared to other genes (median log_2_FC -0.21 vs. −0.07; p=1.2×10⁻^4^, Mann-Whitney test). (P) Expression by lung epithelial cell types in a human IPF scRNA-seq dataset(64) of the 50 genes with the greatest loss of H3K4me3 in response to TGF-β, as determined by H3K4me3 CUT&RUN. (Q,R) Expression of indicated genes, as measured by RNA-seq in iAT2 cells treated with TGF-β in the presence or absence of MI-503. Mean ± SEM. *****padj<0.00001, DESeq2 Wald test with Benjamini–Hochberg correction. **Statistics.** (A,D) One-sample t-test. **(B,C,G)** Ratio paired t-test (log_2_ FC), comparing conditions with versus without SB431542 (B), CHIR and KGF (C), or SIS3 (G), with Benjamini, Krieger and Yekutieli correction. (A,C,D,Q,R) *q < 0.05, **q < 0.01, ***q < 0.001, ****q < 0.0001. (B,G) Statistics in Table S5. n = 3 independent experiments. Error bars: mean ± SEM.

**Supplemental Figure 5.**
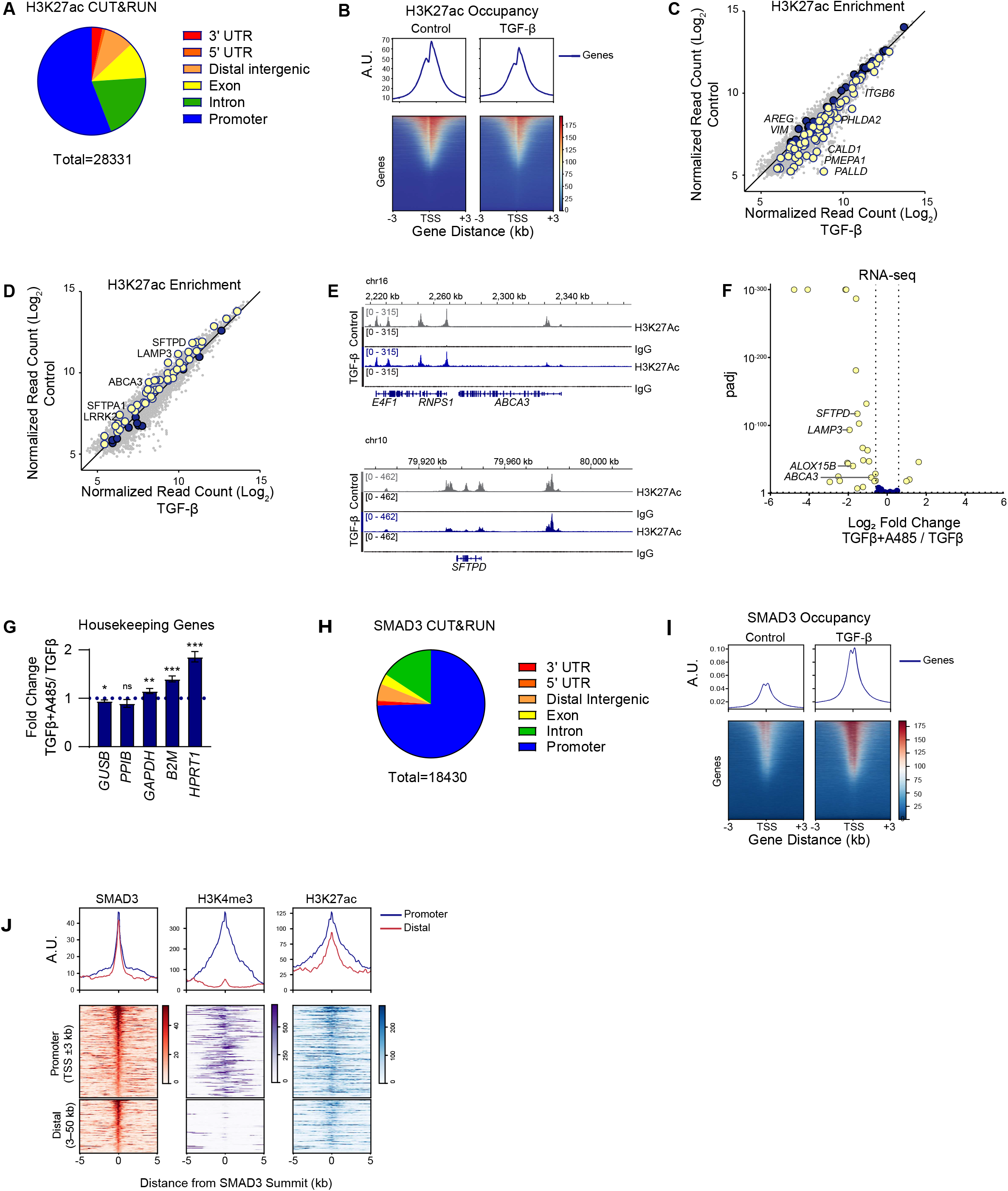
p300/CBP is Required for the AT2-to-*KRT5−/KRT17+* State Transition. (A-E) iAT2 cells treated with TGF-β were subjected to CUT&RUN with an H3K27ac antibody or IgG. (A) Genomic distribution of all called H3K27ac peaks across both control and TGF-β conditions, average of n=3. (B) Genome-wide H3K27ac occupancy shown as TSS-centered profile plots and heatmaps (±3 kb) comparing untreated iAT2 cells (Control) and TGF-β-treated (*KRT5−/KRT17+)* cells. (C) Normalized H3K27ac CUT&RUN read counts (log_2_) at all annotated peaks. Large dots: *KRT5−/KRT17+* markers. *KRT5−/KRT17+* genes with increased (yellow dots) or decreased (blue dots) H3K27ac following TGF-β treatment. No FC or p-value cutoff applied. *KRT5−/KRT17+* marker genes showed gene-set-level H3K27ac gain following TGF-β compared to all other genes (median log_2_FC 0.36 vs. 0.10; p = 1.4×10⁻^6^, Mann-Whitney test). 83.6% of *KRT5−/KRT17+* genes were marked by H3K27ac in untreated iAT2 cells (OR = 2.61 relative to other protein-coding genes; P = 1.1 × 10⁻⁷ by Fisher exact test). (D) Normalized H3K27ac CUT&RUN read counts (log_2_) at all annotated peaks. Large dots: AT2 markers compared to *KRT5−/KRT17+* state(64) with decreased (yellow dots) or increased (blue dots) H3K27ac following TGF-β treatment. No FC or p-value cutoff applied. AT2 marker genes showed a gene-set-level loss of H3K27ac following TGF-β treatment compared to all other expressed genes (median log_2_FC -0.32 vs. -0.10 p=1.9 x 10^-6^, Mann-Whitney test). (E) Representative genome browser tracks showing mean H3K27ac and IgG CUT&RUN signal at the indicated gene loci in untreated (Control) and TGF-β-treated (TGF-β) cells. (F) RNA-seq of iAT2 cells treated with TGF-β ± A485. AT2 genes shown. Yellow dots: |log_2_FC| > 0.59, padj < 0.05. Blue dots: |log_2_FC| ≤ 0.59 or padj ≥ 0.05. Dotted lines at |log₂ FC| = 0.59 and padj = 0.05. (G) Expression of indicated genes by RNA-seq of iAT2 cells treated with TGF-β ± A485. Dotted line: FC = 1. Mean ± SEM. *padj < 0.05, **padj < 0.01, ***padj < 0.001 by DESeq2 Wald test, Benjamini-Hochberg adjusted. (H) Genomic distribution of all called SMAD3 peaks across both control and TGF-β conditions, average of n=2. (I) Genome-wide SMAD3 occupancy shown as TSS-centered profile plots and heatmaps (±3 kb) comparing untreated iAT2 cells (Control) and TGF-β-treated (*KRT5−/KRT17+)* cells. (J) Heatmaps of SMAD3, H3K4me3, and H3K27ac CUT&RUN signal in TGF-β–treated iAT2 cells, centered on SMAD3 peak summits (±5 kb). Rows represent SMAD3 peaks within 50 kb of the transcription start sites (TSSs) of *KRT5−/KRT17+* genes, grouped as promoter (TSS ±3 kb; n = 209) or distal (3–50 kb from the TSS; n = 121) and sorted within each group by SMAD3 signal at the summit. Average signal profiles for promoter (blue) and distal (red) peaks are shown above each heatmap. A.U.: log_2_ fold over IgG for SMAD3; TMM-normalized coverage for H3K4me3 and H3K27ac. n=2. (A-K) n=3 independent experiments. (A-E) Differential binding was assessed by DiffBind/DESeq2 with Benjamini–Hochberg correction.

**Supplemental Fig. 6.**
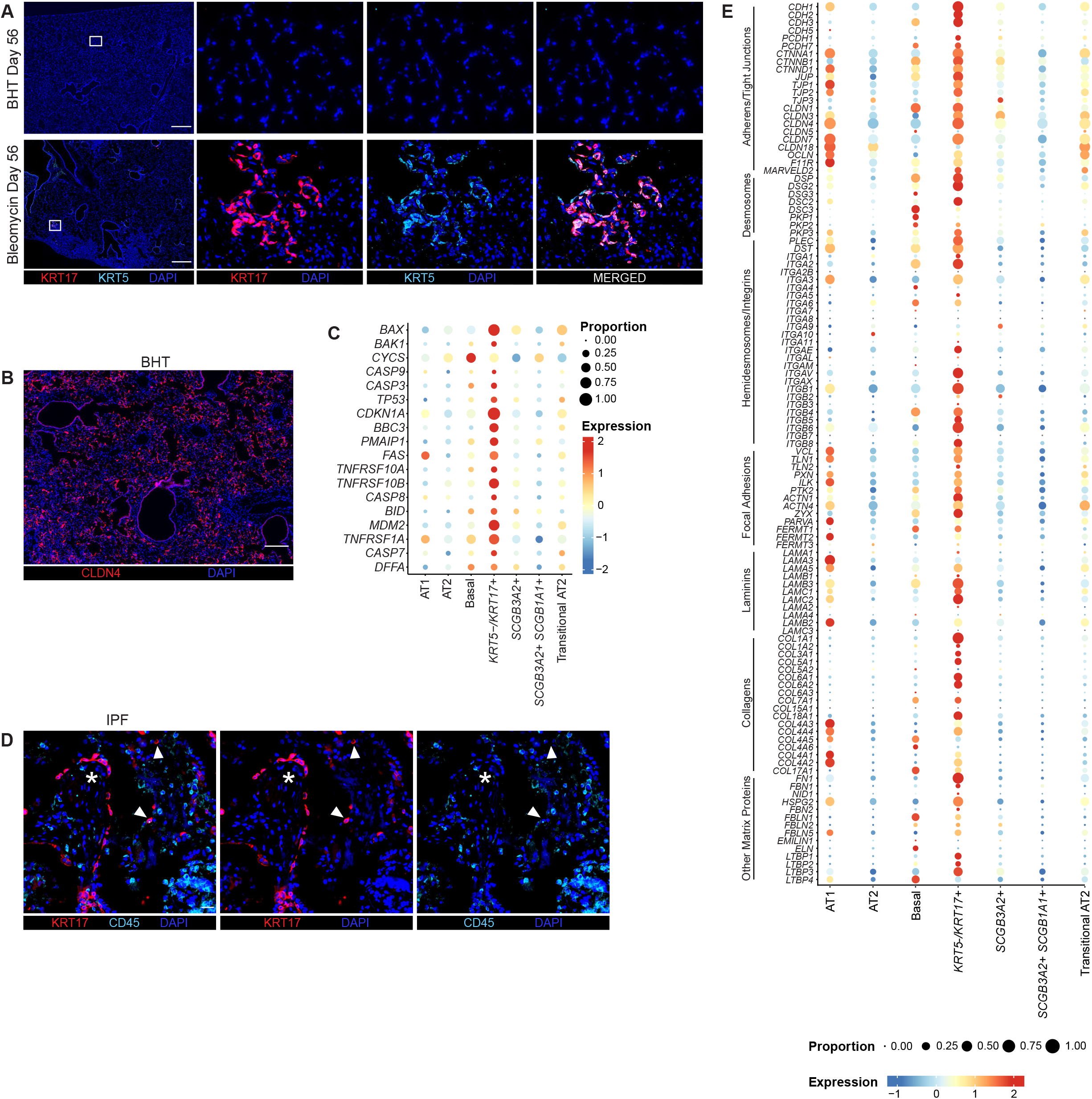
*KRT5−/KRT17+* Cells Undergo Turnover. (A) Staining of murine lung after bleomycin or BHT injury for indicated antibodies. (B) Immunostaining of BHT-treated lungs for indicated antibodies. (C) Expression of apoptosis markers by *KRT5-/KRT17+* cells in IPF, as determined by scRNA-seq(64). (D) Immunostaining of IPF lung for indicated antibodies. (E) Expression of cell-cell adhesion genes by *KRT5-/KRT17+* cells in IPF, as determined by scRNA-seq(64). Scale Bars: 500 µm (A), 200 µm (B), and 50 µm (D). n=3-6 mice/group

**Supplemental Figure 7.**
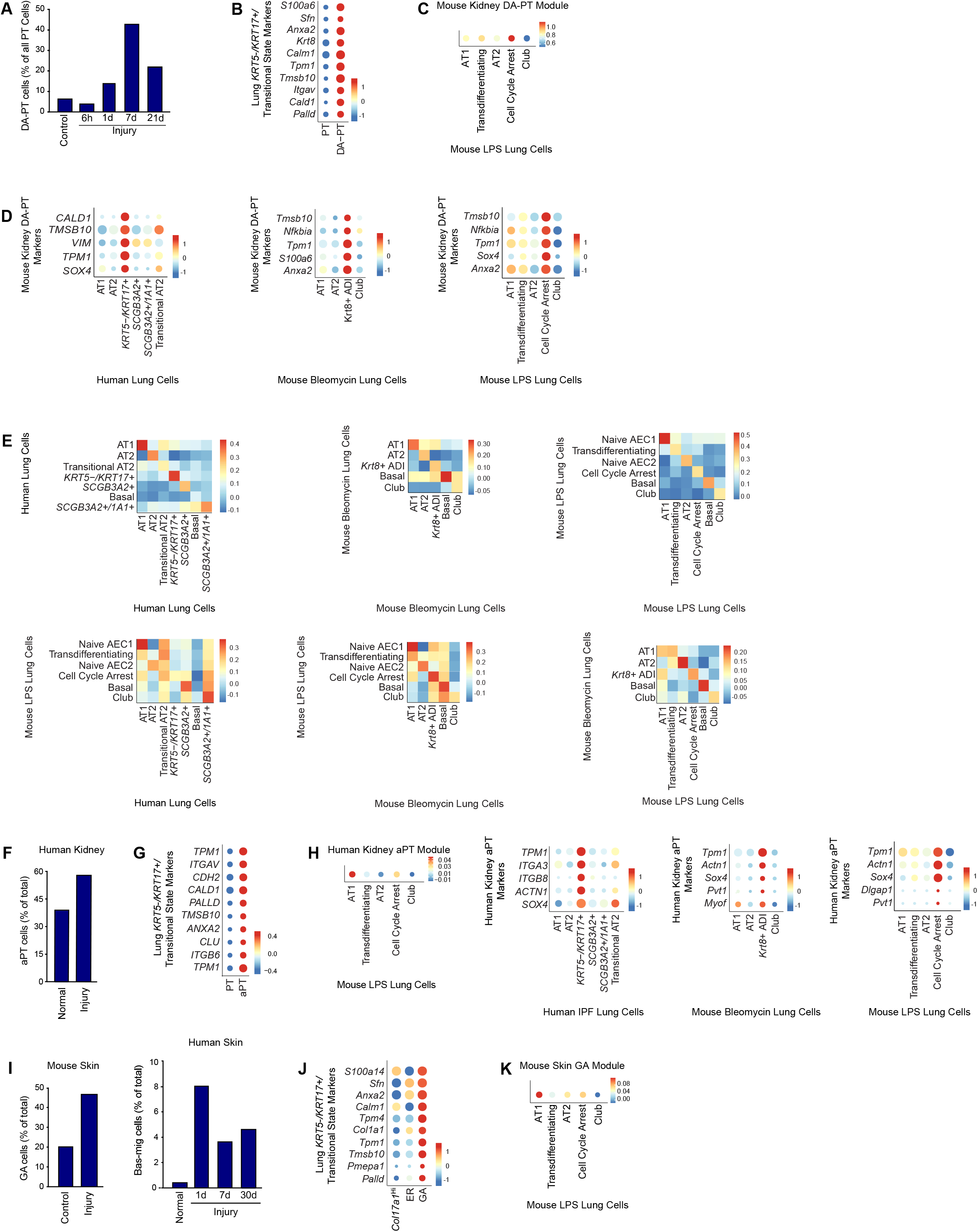
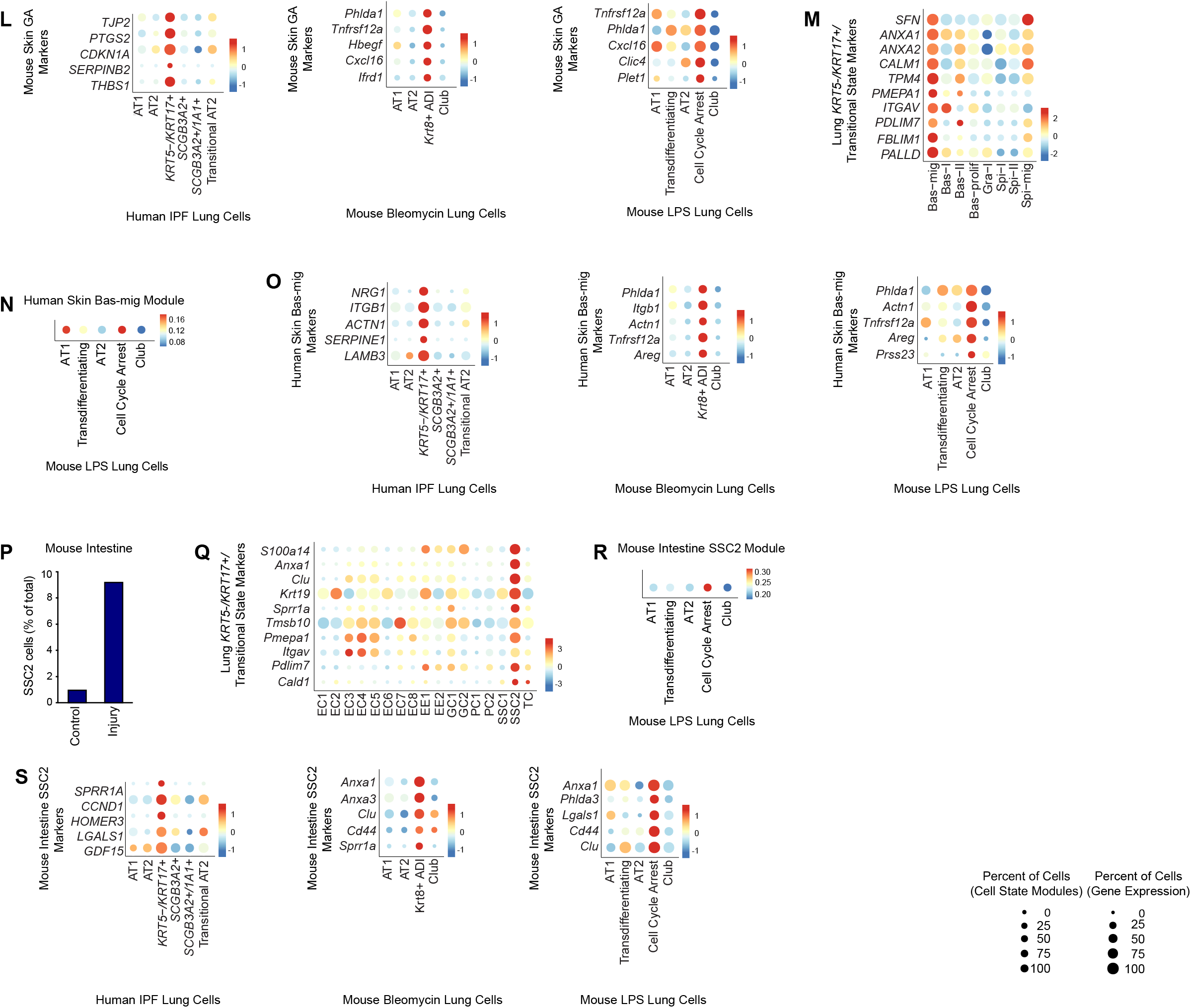
Conserved Cell States Emerge from Lung, Kidney, Skin, and Intestine. (A,F,I,P) Percentage of injury-emergent cells in control and injured tissue: (A) DA-PT cells in mouse kidney(92); (F) aPT cells in human kidney(93); (I) GA basal cells in mouse skin(95); Bas-mig cells in human skin(96); and (P) SSC2 cells in mouse small intestine(94). (B,G,J,M,Q) Expression of lung *KRT5−/KRT17+*/transitional state marker genes by (B) mouse kidney PT, (G) human kidney PT, (J) mouse skin, (M) human skin, and (Q) mouse small intestine epithelial cell states. (C,H,K,N,R) Gene module activation of the (C) mouse kidney DA-PT, (H) human kidney aPT, (K) mouse skin GA, (N) human skin Bas-mig, and (R) mouse intestine SSC2 states by mouse lung epithelial cell types(80). Dot size represents percent of cells with a Cell State Module score > 0. (D,H,L,O,S) Expression of (D) mouse kidney DA-PT, (H) human kidney aPT, (L) mouse skin GA, (O) human skin Bas-mig, and (S) mouse intestine SSC2 state marker genes by human and mouse lung cell types(63, 64, 80). (E) Mean scTOP score for each lung cell type compared with reference lung cell types in all pairwise query/reference combinations(63, 64, 80). Reference lung cell types on the x-axis. Scale bars indicate the mean scTOP score. (B,D,G,J,L,M,O,Q,S) Dot size represents the percent of cells with nonzero expression of the indicated gene. Scale bars indicate average gene expression as a z-score. ADI, alveolar differentiation intermediate; Bas, basal; Bas-mig, basal-migrating; Bas-prolif, basal-proliferating; Col17a1Hi, collagen type XVII alpha 1 chain; EC, enterocyte; EE, enteroendocrine cell; ER, early-response; GA, growth-arrested; GC, goblet cell; Gra, granulocyte; PC, Paneth cell; Spi, spinous; Spi-mig, spinous-migrating; SSC, singly profiled stem cell cluster; TC, tuft cell.

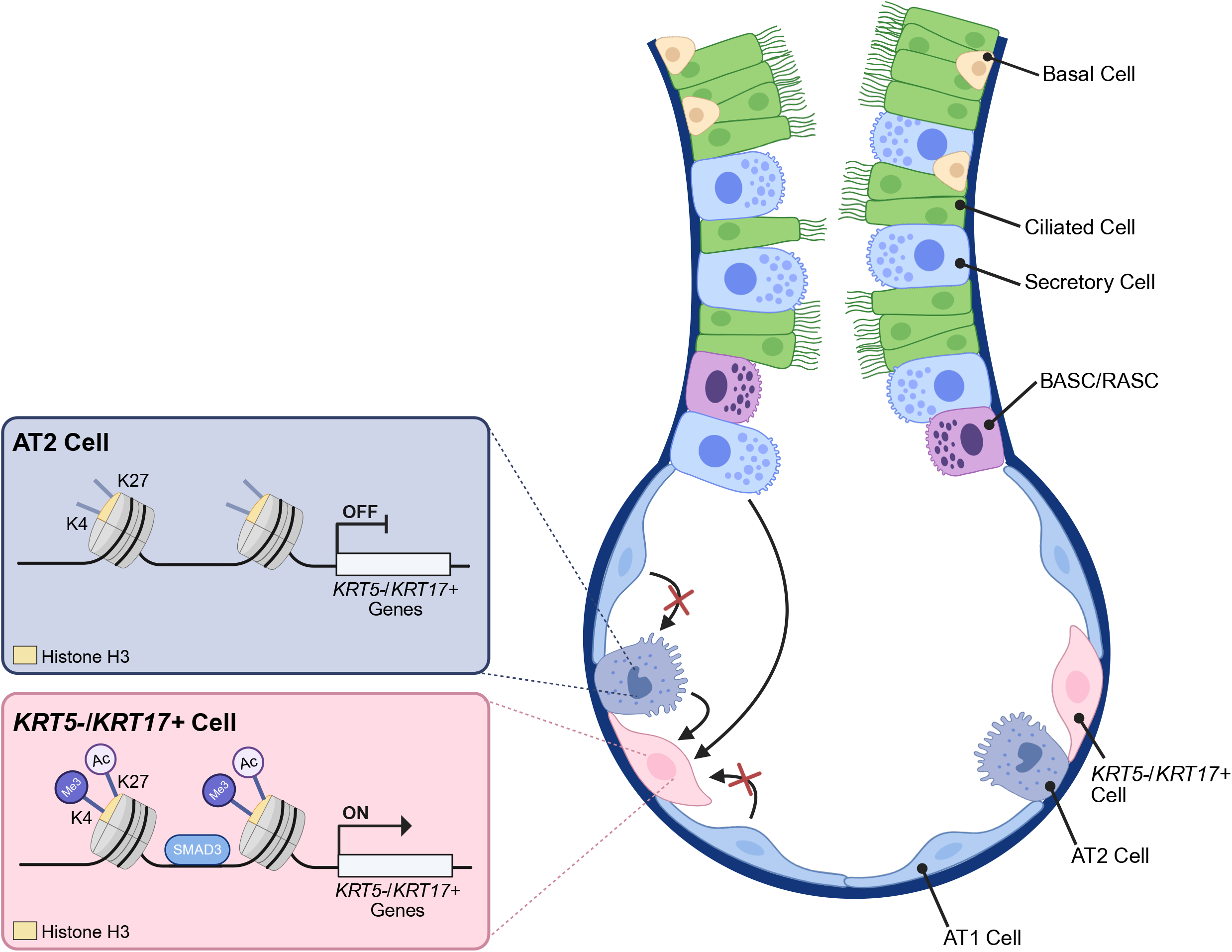

